# Better together – relative retention time plus spectral matching improves automated glycan characterization using PGC-nLC-IT-ESI-MS/MS

**DOI:** 10.1101/2021.08.04.451140

**Authors:** Kathirvel Alagesan, Falko Schirmeister, Uwe Möginger, Arun Everest-Dass, Friedrich Altmann, Peter H Seeberger, Mark von Itzstein, Nicolle H Packer, Daniel Kolarich

**Author notes:** **Corresponding Authors** Dr. Kathirvel Alagesan, ***Present address:*** Max Planck Unit for the Science of Pathogens, Charitéplatz 1, 10117 Berlin, Germany, A/Prof. Daniel Kolarich, Institute for Glycomics, Griffith University, Gold Coast campus, QLD 4222. Novo Nordisk, Novo Nordisk Park 2760 Måløv, Denmark (Uwe Möginger). **CREDIT AUTHOR STATEMENT**. Kathirvel Alagesan – Conceptualization, Methodology, Software, Validation, Investigation, Formal analysis, data curation, Writing – Original draft, Writing – Review & Editing, Visualization, Project administration. Falko Schirmeister – Investigation, Software, Validation, Writing – Review & Editing, Visualization. Uwe Möginger – Investigation, Validation, Writing – Review & Editing, Visualization. Arun Everest-Dass - Validation, Writing – Review & Editing, Visualization. Friedrich Altmann – Resources, Writing – Review & Editing. Peter H Seeberger – Resources, Writing – Review & Editing. Mark von Itzstein – Resources, Writing – Review & Editing. Nicolle H Packer – Resources, Writing – Review & Editing. Daniel Kolarich – Conceptualization, Methodology, Resources, Visualization, Writing – Review & Editing, Funding acquisition.

## Abstract

Porous Graphitized Carbon nano-liquid chromatography tandem mass spectrometry (PGC-nLC-MS/MS) is a glycomics technique with the unique capacity to differentiate isobaric glycans. The lack of suitable software tools integrating chromatography and MS-information delivered by PGC-nLC-MS/MS has been limiting fast and robust glycan identification and quantitation. We report a LC-system-independent strategy called GlycoRRT that combines relative retention time (RRT) and negative ion fragment spectra analyses for isobaric structure-specific glycomics of PGC-nLC-MS/MS data. The GlycoRRT toolset is fully customizable and easily adaptable enabling semi-automated high-throughput structural assignments. The current library contains over 200 entries and their individual meta-data (MS instrumentation, experimental conditions, retention times, fragmentation profiles and glycan structural diagnostic ion features) relevant for reliable data analyses. The GlycoRRT workflow was employed to map the N*-* and O*-*glycome in blood group matched human plasma and urine as well as decipher Immunoglobulin (IgG) glycosylation features from 13 different animal species. We have also developed visualization tools to enable a consistent, reliable, and reproducible analysis of large sets of multidimensional PGC-nLC-MS/MS glycomics data. This comprehensive glycan resource provides the glycan map of human and animal species, will serve as a reference in dissecting the role of glycans in host pathogen interaction and zoonotic disease transmission.

---

Cell surface and body fluid proteins are extensively modified with specific sugar moieties, so-called glycans ^1^. These glycans build the basis for a universal language (glycome) that is used between cells but is also exploited by pathogens and cancer cells ^1, 2^. Glycomics seeks to understand how the entirety of glycans present on a protein, a cell or a body fluid relate to biological processes in health and disease by detailed structural and semi-quantitative analyses of glycans ^3, 4^.

In order to identify and quantify the cells’ complete glycan repertoire, it is necessary to capture the enormous structural diversity and complexity. This daunting task requires orthogonal methods to accommodate both, in-depth structural data acquisition and medium-high throughput analyses ^5^. Widely used Mass Spectrometry (MS) or Liquid Chromatography (LC)-based approaches provide good throughput but suffer from limited capacity to elucidate structure details and differentiate isobaric glycans. In consequence, the combination of LC separation with tandem mass spectrometry (MS/MS) has evolved to be one of the most powerful platforms to discriminate, distinguish, and quantify glycan structural isomers ^3, 4, 6^. Separation of structure isomers has been achieved by reversed-phase (RP) and hydrophilic liquid interaction chromatography (HILIC) ^7, 8^, but porous graphitized carbon (PGC) presents itself as the most versatile and effective approach to separate isobaric glycans ^6, 9, 10^. PGC can be applied universally to analyze not only N*-* and O-glycans, but also the glycan portion of glycolipids and glycosaminoglycan fragments ^11-13^. In negative ion mode, PGC-nLC-MS/MS allows for an unambiguous and detailed structural characterization of isobaric glycans and relative quantitation within a single LC-MS/MS experiment, drastically reducing the sample amount to even capture the glycome from as few as 1000 cells ^14-17^.

Unlike HILIC, glycan retention in PGC cannot be correlated to the monosaccharide composition by multiple linear regression analyses ^18^. The exact molecular nature of the PGC-retention mechanisms leading to the observed isobaric glycan separation is still not fully understood, further complicating computationally assisted retention time predictions. Nevertheless, specific glycan structural features are known to influence the PGC elution behavior, and the order of elution of individual structural isomers is also highly conserved ^19-21^. Despite the availability of glycan retention time and elution order, efforts to utilize them for glycan identification have not yet been satisfactorily implemented as the absolute retention times can vary between individual analyses and drastically depend on the individual LC-setup (e.g. capillary lengths) and operational parameters (organic phase gradient) used.

Absolute retention time variations can be compensated by normalizing retention times based on calibration standards. This approach has been implemented successfully for the automated structure assignment of HILIC-separated 2-AB-labeled glycans. The detected retention times are converted into “Glucose Units” (GU) determined by a separate analysis of a dextran ladder ranging from 1-15 glucose molecules ^22^. A similar normalizing approach has been described recently for PGC-LC-MS/MS where the glycan structure assignment is based only on precursor mass and GU values with subsequent verification by fragment spectra ^23^. Beforehand, Pabst and co-workers were the first to demonstrate the use of relative retention times (RRT) for structural assignment of oligomannosidic N-glycans ^24^. However, this promising development has not been fully utilized for other classes of glycans that can be separated using PGC-LC. This is not surprising, as RRT and precursor MS is insufficient for reliable structure determination considering the tremendous structural diversity represented by glycans ^9, 25^.

To this end, a spectra library-based approach that allows for querying multiple tandem spectra against a well-defined library alongside precursor mass and RRT will improve glycan structure assignment and facilitate high-throughput (HTP) data analyses. Currently available software tools that predict glycan structures from MS data do not allow for a direct comparison of query spectra against library spectra ^26^. We addressed these challenges and developed a system-independent, semi-automated and easily adaptable toolset for automated glycan identification that combines MS/MS spectra matching, normalized glycan elution order and precursor mass. These distinct parameters along with peak intensity are acquired within a single PGC-nLC-MS/MS analysis for each glycan structure.

The herein described GlycoRRT tool incorporates and fully utilizes these distinct features to allow for glycan structure assignment and refinement as well as relative quantitation. The initial GlycoRRT library was established using data of 33 synthetic N-glycan standards and has been subsequently extended to now contain 200 annotated N-glycans from human plasma glycoproteins. Using the GlycoRRT tool, we established the human GlycoATLAS, a new resource that provides a baseline map of the N*-*glycans of human whole blood as well as blood group matched pooled plasma and urine. Finally, we also mapped species specific immunoglobulin (IgG) glycosylation features of 13 different animal species.

## RESULTS

### CREATION OF GLYCORRT SPECTRAL LIBRARY

A reliable, fully customizable GlycoRRT spectral library was developed based on PGC-nLC-ESI-MS/MS in negative ion mode to enable HTP automated glycan identification and characterization. The initial GlycoRRT spectral library was established utilizing 33 synthetic N-glycan standards (**Table S1**) incorporating two distinctive features: a) high-quality negative ion fragment spectra (**Figure S1&2**) and b) retention time (**Table S3**). In addition to the expected 33 N-glycans, we were also able to resolve other isomers present in these standards yielding a total of 41 distinctive tandem MS spectra (**Table S3)**. Each entry in the GlycoRRT spectral library contained all crucial metadata including precursor mass, charge state, glycan composition, and structure represented in GlycoMod and Proglycan nomenclature ^27^ (**Figure S3**). An overview of the GlycoRRT workflow is provided in **Figure 1** and described in the supplementary methods section.

**Figure 1:**
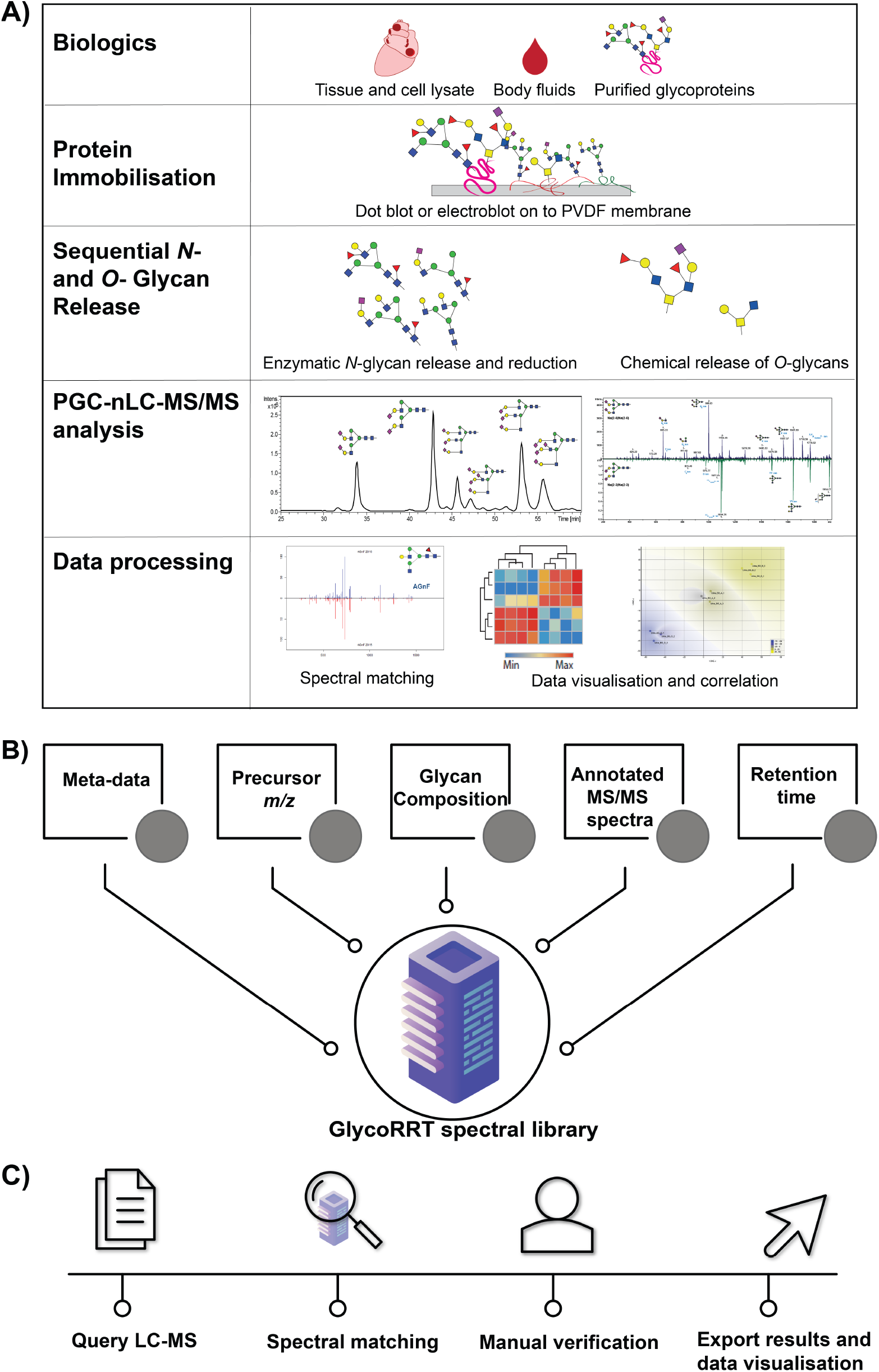
PGC-LC-ESI-MSMS and GlycoRRT spectral-library matching glycomics workflow. **(A)** The four main steps of glycomics sample preparation: The proteins were extracted from the desired biological material using suitable methods and then immobilized onto a PVDF membrane. N-linked glycans were released enzymatically using PNGase F, followed by the O-glycan release from the same sample by reductive β-elimination. Finally, N- and O-glycan alditols were desalted and analyzed by PGC-nLC-ESI MSMS. **(B)** Glycan spectral library design: The Glycan RRT LC–MS/MS library contains annotated, negative ion ESI-MS product ion spectra, precursor ion information including charge state, retention time information, glycan composition and isomer information, as well experimental metadata. **(C)** Spectral matching of query spectra against the well-annotated reference spectra database enables automated glycan identification, providing as %fit score for each glycan that informs on positive or negative identification. The results can be easily exported and are readily visualized using our custom R packages.

#### Spectral matching evaluation

We demonstrated previously that it is possible to use the spectral matching approach for automated glycan structure characterization based on the dot-product function available in the R package OrgMassSpecR ^28^. Nevertheless, the spectral matching function was not available within the data analysis workflow for HTP analyses. We first evaluated the feasibility of the spectral matching tool incorporated within the data analysis software by re-searching the dataset used to create the initial spectral library. The query compounds were identified in high confidence as evident by the high RFit’ score (max 1000). Next, the applicability and feasibility of the spectral library to identify N-glycans derived from a randomly selected human Immunoglobulin (IgG3) was evaluated. Details on the workflow for the sequential release of N- and O-glycans adapted from a previous protocol ^10^ are described in the online methods. The initial analyses resulted in the identification of 15 different N*-*glycan structures from 500 ng protein in less than minute via single click data analysis **(Table S4, Supplementary results - Spectral matching automation in data analysis)**. The RFit’ scores of the identified N-glycans ranged from 918 for GnA to 988 for Na(2-6)Na(2-6) indicating high confidence in the structure assignment (**Table S4**). Nevertheless, the entire data file contained several spectra that did not result in any confident library match at that time point. The following manual analyses focused on these unidentified spectra, adding another six N-glycans to the spectra library. We confidently identified 21 different N-glycan structures from 500 ng human IgG3 in less than 15 minutes using this automated spectral matching and manual identification approach (***Table S6 – for additional identifications***).

#### Relative Retention time inclusion

Spectral matching provides an enormous step forward in automated structure identification; however, closely related, isobaric glycans often share multiple product ion spectra features that can result in incorrect structure assignments. Such wrong assignments can be reduced significantly when glycan elution profile information is included ^11, 19^ (***Figure S4***). Thus, the integration of glycan structure isomer information derived from the relative order of PGC LC-elution together with spectral matching can facilitate and improve HTP automated glycan structure assignment.

Run-to-run variation in the absolute retention time and among various laboratories, however, impedes the integration of absolute retention time information that will assist in the structural assignment. We first evaluated the influence of column temperature and grounding on the separation capacity offered by PGC to better understand the impact of these conditions on PGC separation in our LC-MS setup (**Supplementary Results: Glycan elution Rules of PGC & Factors influencing glycan separation by PGC-LC, Figure S5 & S6**) ^29^. Next, we sought to normalize the variation in absolute retention times by converting these values into relative retention times (RRT) and incorporated this feature into our spectral matching tool. The RRT for each glycan species was calculated using the AAF (***N-glycan ID20, Table S5***) isomer elution time as a reference value, as it elutes in the middle of the gradient. The RRT values of all other synthetic N-glycan standards were then established in relation to AAF (***Table S5***). The robustness of RRT normalization was then evaluated using 12 independent samples obtained from three different users containing 21 N-glycan species that were all identified by using the spectral matching tool as described above with high accuracy (**Table S6, Figure 2A**). The RRT approach corrected absolute retention time shifts with less than 5% coefficient of variation (**Figure 2-B**).

**Figure 2:**
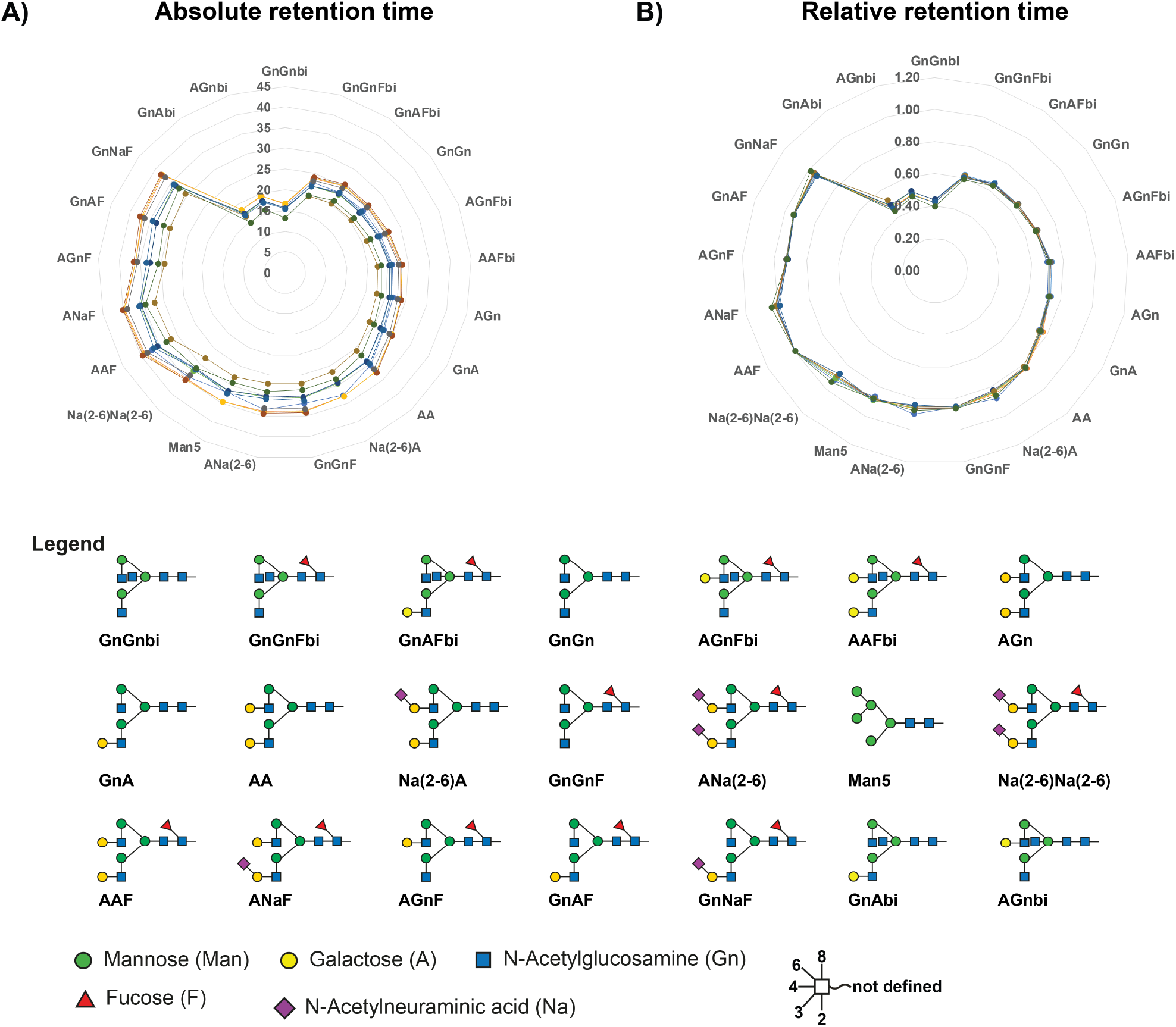
**(A)** Absolute retention times of 21 N-glycans collected from 12 independent samples that were obtained from three different users and their relative retention times (RRT). **(B)** The RRT for each identified N-glycan was calculated using the AAF isomer elution time as reference using the expression RRT = (Sample RT / Reference RT). The use of RRT normalizes the variations in absolute retention time for accurate incorporation of retention time features in glycan structure assignment. Glycans are labeled with their simple “Proglycan” nomenclature to allows for differentiation of glycan isomers.

We also evaluated the influence of the ion trap instrument’s Smart Parameter Setting (SPS) target parameter on glycan identification and, specifically, quantitation. Typically, the SPS is set in the middle of the intended acquisition mass range to support the auto-adjustment of the specific acquisition mass window to a particular target *m/z*. Our results indicate that the selection of an SPS target *m/z* oriented slightly towards the higher end of the intended *m/z* scan range improved the identification and quantitation of larger glycan structures compared to the traditional mid-range SPS selection (***Figure S7, see supplementary results - Influence of SPS parameter on quantification***).

### THE HUMAN GLYCOATLAS

The results of the initial GlycoRRT approach for HTP glycan analyses motivated us to further expand the library using N-glycans released from human serum glycoproteins.

#### Plasma/serum vs whole blood N-glycan atlas

The serum GlycoRRT spectral library was used to profile plasma and whole blood N-glycans. In total, 70 different compounds corresponding to N-glycans were identified in less than 5 min, including various charge states (**Table S7, Figure S8-A**). In addition to the RFit’ score, relative retention times as well as any deviations (dRRT, ***Formula: dRRT = RRT library – RRT Run***) were used as a means of quality control for unambiguous glycan structure identification and characterization. Of 70 N-glycan compounds identified (60 by automated identification, two via manual identification, the remaining eight are different charge state variants of glycans already present in the library), the relative abundances of 68 N-glycan structures were quantified by integrating the area under the curve obtained for the respective EICs. For two tetra-sialylated N-glycans the EIC-peak was unsuitable for reliable quantitation. For each measurement, the values were normalized to the total quantity of all detected N-glycans.

Next, we sought to visualize the difference in relative abundance at both individual glycan and group level. The relative abundances of individual N-glycan structures were assessed and visualized in a heatmap by unsupervised hierarchical cluster analysis **Figure S8-A (See Table S7 for identification results)**. The most abundant N-glycan in serum, plasma, and whole blood was the Na(2-6)Na(2-6) glycan contributing to approximately 50% in all three types of samples (**Figure S8-A**). The N-glycans were also grouped based on their structural features: Oligomannose, neutral complex and sialylated complex. Sialylated N-glycans accounted for approximately 90% of the total N-glycome in the analyzed body fluids. The majority of these were di-sialylated (75% in the case of plasma and serum, 70% in whole blood), followed by mono- and tri-sialylated. Tetra-sialylated N-glycans were just present in trace amounts (**Figure S8-B**). Overall, the N-glycans derived from plasma and serum exhibited similar relative abundances for all glycan classes, whereas serum/plasma vs whole blood exhibited marked differences especially in the case of mono- and di-sialylated N-glycans (**Figure S8-B**).

#### Blood-group specific features of the plasma and urinary N-glycome

The GlycoRRT tool was further employed to establish an average profile for the global plasma and urine N-glycome (600 ng - injected) after pooling these samples according to blood groups (A, B and O). After automated and manual identification, 46 N-glycan isomers corresponding to 29 glycan compositions were detected in plasma, whereas 33 N-glycan isomers corresponding to 20 glycan compositions were identified in urine by a single click analysis that took less than 15 min analysis time for 18 samples on a standard PC (**Table S8 and S9)**. The N-glycan structures initially not present in the library such as tri-antennary N-glycans from plasma (Glycan ID: 32 and 33 – **see Table S9**) and N-glycans carrying the LacdiNAc epitope (Glycan ID: U23 and U24, see Table S11) from urine were manually identified, validated, added to the library, and quantified and visualized in a heatmap (**Figure 3 and 4)**. Na(2-6)Na(2-6) was the most abundant glycan in both plasma and urine irrespective of the blood group.

**Figure 3:**
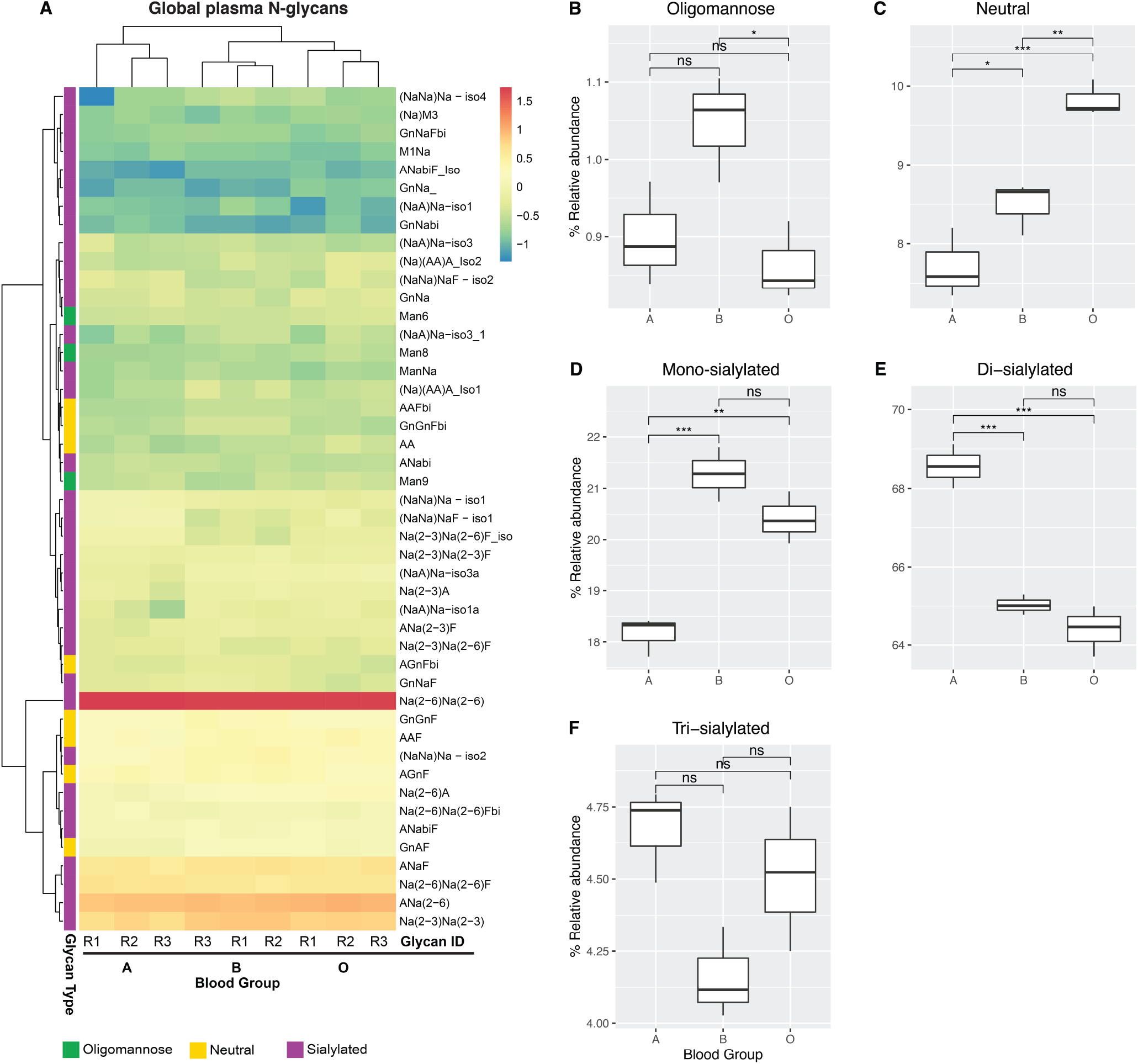
(A) Heatmap of log-transformed relative abundances of N-glycans from blood group matched plasma. Rows indicate the glycan structures whereas the column indicate the three technical replicates. from different blood group (A, B and O). The samples were ordered using unsupervised hierarchical clustering based the Euclidean distance matrix with complete linkage. Remarkably, the overall glycome signature allowed the algorithm to independently differentiate the pooled plasma samples based on their blood groups indicating a blood group dependent impact on the global N-glycome profile. (B-F) Differences in relative abundance of different N-glycan classes are shown as box plots. Multiple comparisons among various blood groups were performed using paired T-test. The segregation of blood group B and O from A is largely driven by differences in the abundance of mono and di-sialylated N-glycans. See source data table S1 for pair-wise comparison and results table.

**Figure 4:**
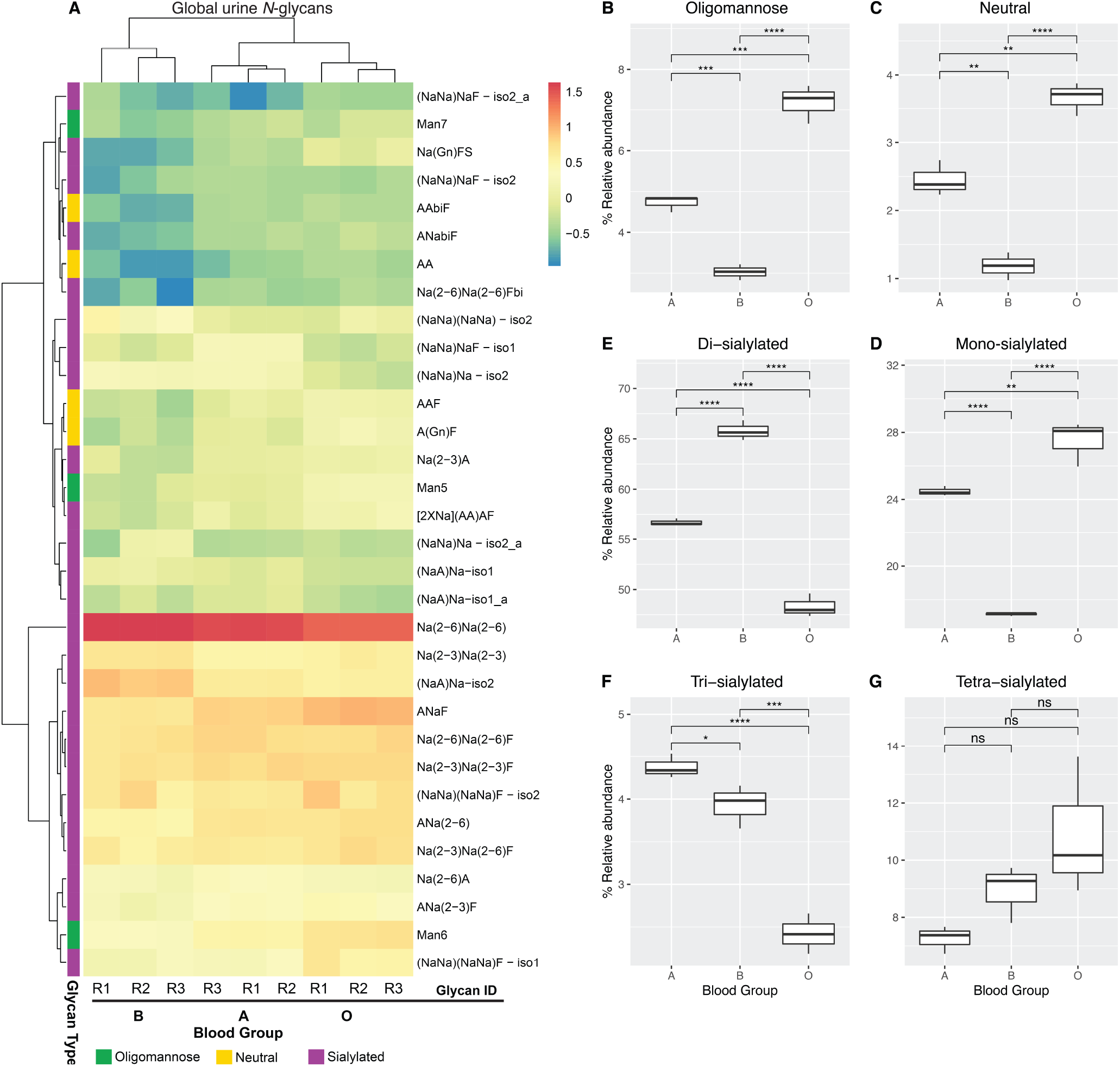
(A) Heatmap of log-transformed relative abundances of N-glycans from blood group matched urine. The samples were ordered by hierarchical clustering using the Euclidean distance matrix with complete linkage. Rows indicate the glycan structures (Supplementary Table S9). Similar to plasma glycome, also the urinary N-glycome enabled the algorithm to cluster the samples based on the ABO blood group status. We also observed two clusters based on unsupervised hierarchical clustering yet in an opposite trend compared to plasma indicating source dependent differential action of glycan modifying enzymes. (B-F) Differences in relative abundances of different N-glycan classes are shown as box plots. Multiple comparisons among various blood groups were performed using paired T-test. As evident, the segregation of N-glycans derived from blood group B and O from A is largely driven by mono and di-sialylated N-glycans. See source data table S2 for pair-wise comparison and results table.

#### A snapshot of Vertebrata Immunoglobulin G (IgG) glycosylation

The developed PGC-nLC-ESI-MS/MS glycomics workflow was also utilized to investigate IgG N-glycans derived from thirteen different Vertebrata species (**Table S10**). Samples from twelve mammalian and one avian species were compared with regard to the similarity of their qualitative and relative quantitative IgG N-glycan signatures (**Figure S9 & Figure 5**). The hierarchical cluster analysis sorted the animals in an unbiased manner, solely based on the determined N-glycan profiles. Most species showed a preference for incorporation of either NeuAc or NeuGc. The human IgG N-glycans separated clearly from the other mammals due to the lack of N-glycans carrying NeuGc. Duck IgG N-glycans showed the highest degree of differentiation due to the presence of unique N-glycan structures exhibiting sialylated LacdiNAc motifs not found in any other species analyzed (**Figure S9**). Vertebrate IgG contained predominantly contained biantennary neutral N*-*glycan followed by sialylated N-glycan except in the case of Donkey (**Figure 5 A-C**).

**Figure 5.**
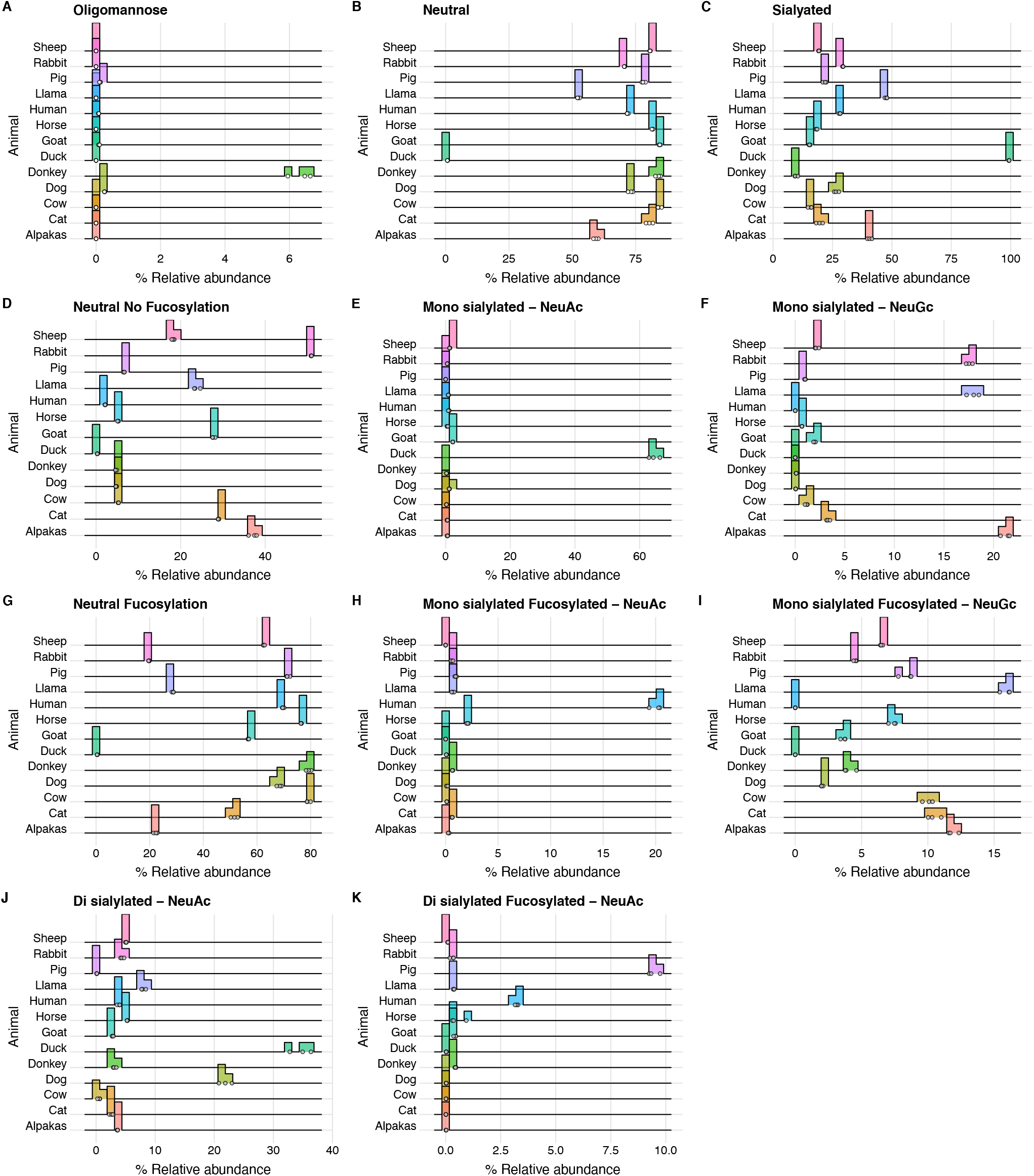
**A-K:** Rigdeline plot of Glyco-epitope features of IgG N-glycan derived from twelve different mammals and one bird. Glyco-epitope features detected exhibited a strong species dependency. Irrespective of the fucosylation status, the bulk of the IgG N-glycans were neutral ones lacking NeuAc or NeuGc, with the exception of donkey who predominately contained sialylated N-glycans (**A-C**). The majority of the neutral N-glycans were core-fucosylated except for duck who was lacking fucosylation on its IgG (**D**,**G**). Incorporation of either NeuAc or NeuGc was dependent on the species. Human and Duck contained only NeuAc. Nevertheless, the NeuAc present in duck was just found attached to a LacdiNAc motif which was not found in any other species. Human and dog had similar NeuAc levels but exhibited different levels of core fucosylation (**E-K**).

## DISCUSSION

Despite the unique analytical opportunities provided by various emerging LC-MS technologies to characterize glycans, reliable interpretation of repetitive glycan MS/MS data is a major limiting factor in HTP data analysis. **Figure 1-B&C** summarizes our GlycoRRT pipeline for HTP qualitative and quantitative assessment of multi-dimensional glycomics data obtained using PGC-nLC-ESI-MSMS in negative ion mode. The conserved diagnostic fragment ions observed in product ion spectra of deprotonated glycan ions ^30-32^ in combination with the high separation capacity provided by the PGC was the basis to develop this reliable, fully customizable GlycoRRT tool. To best of our knowledge, this study represents the first examination towards optimizing and improving the existing tool for HTP glycan structural assignment. Our GlycoRRT tool utilizes (i) retention time and glycan elution order, (ii) precursor mass, and (iii) fragment spectra to achieve reliable structure identification.

Current bioinformatics tools for MS/MS-based glycan sequencing rely on inferring glycan composition from the precursor mass using tools such as GlycoMod ^33^ and use information derived from diagnostic fragment ions in the product ion spectra for structure determination. Despite the availability of the Glycoworkbench ^34^ software that assists in manual interpretation of tandem MS data, the process is tedious and repetitive. Spectral library matching for N-glycans has been reported previously to show good spectral reproducibility and glycan fragmentation in negative ion mode and is fully suitable for facilitating cross-platform analyses ^28^. To this end, a spectral matching approach including standardized retention time data for structure assignment enables reliable HTP data analysis while minimizing repetitive tasks. Retention time information is regularly used in normal phase separation for glycan identification, but it is less frequently an integrative parameter in LC-MS/MS based glycomics to support structure identification and/or additional validation. However, the absolute retention time of glycan analytes can vary substantially when samples are analyzed across a long time span as it is required when investigating large cohorts. Here we relied on relative retention times to normalize the unavoidable variation in the absolute retention time. We show that the RRTs are system independent and remain stable over an extended time period for years (**Figure 2A**). RRT values are calculated by dividing each individual retention time value by the retention time of one selected reference glycan structure (we selected AAF for our analyses). In principle, however, any glycan structure and its corresponding retention time can be used as reference as long as they are present in all the samples and as such, the proposed system is highly flexible and can be optimized for any sample cohort.

We created the GlycoRRT tool using the LibraryEditor available within the DataAnalysis software (Bruker) that allows creation of glycan spectra data bases as well as fast spectra matching to facilitate a fast and reliable, semi-automated data analysis. Nonetheless, any available spectral matching tool such as MSdial ^35^ can be refined for automated glycan identification by incorporating retention time, associated experimental, MS and MS/MS acquisition metadata. Caveats of any software-assisted approach relying on spectral library matching are false assignments or low scores, especially if fragment ions are of low intensity, absent or very limited (as it is frequently found for highly sialylated glycan precursors). However, the approach presented here enables easy identification of such wrong or no assignments based on RFit score and dRRT values. Upon manual validation, the new structures can be easily added as a new entry to the library. Including GlycoRRT also allows identifying highly sialylated N-glycans that deliver poor product ion spectra with improved accuracy. Overall, our work represents a first step towards automated qualitative glycan analysis by incorporating spectral matching and RRT.

Relative quantitation allows for the sensitive monitoring of changes in glycan abundances across various samples and biological conditions ^36^. Here, we present several ways to analyze and visualize large-scale multi-dimensional LC-MS glycomics data as a part of the GlycoRRT glycomics workflow. The developed R package *BCQA* covers the data handling from import of raw quantitation values from Bruker’s QuantAnalysis, their normalization, statistical analysis and comprehensive data visualization (**Figure S11**). Furthermore, the developed functions allow seamless conversion of glycan composition to other formats to allow for visualization using other tools available in Glycomics ExPASy server (https://www.expasy.org/glycomics, **See supplementary Figure S12**).

In line with recent global initiatives (https://human-glycome.org/), the developed PGC glycomics workflow was used to capture an overview of the human plasma and urinary N-glycome from blood-group matched plasma and urine pooled from 20 individuals (**Figure 3&4**). Our analyses have demonstrated that the plasma and urinary N-glycome exhibited an ABO blood-group dependent profile (**Figure 3&4**). It has been reported previously that sialylated glycan clusters on the red blood cell membrane are unique for each blood type ^37^. We observed a similar inclination on the secreted glycoproteins present in plasma with a significant increase in di-sialylated N-glycans in plasma from blood group A individuals, while monosialylated were increased in blood group B individuals and blood group O individuals led in neutral N-glycan abundances (**Figure 3 D&E**). In urine, remarkably, blood group B exhibited the highest levels of di-sialylated N-glycans (***Figure 4-E***). To the best of our knowledge plasma glycan biomarker studies have not, included any data stratification based on blood-group status. Our data, however, hint towards the necessity to include blood group stratification and appropriate blood group stratified controls in any type of plasma glyco-marker studies.

Revealing deep N-glycan structure details of species-specific IgGs using our GlycoRRT approach also enabled us to reconstruct the enzymatic steps in N*-*glycan biosynthesis pathways among the 13 different species analyzed (**Figure 6**). Our results clearly indicate that the mechanisms regulating IgG glycosylation are conserved across the different species to deliver predominately biantennary N-glycans (**Figure S10**). The terminal glyco-epitope features, however, are highly species specific (Na in humans vs Na/Ng in other mammals vs LacDiNAc in duck). This is likely the consequence of differences in monosaccharide donor availability, transferase substrate specificities as well as transferase expression.

**Figure 6:**
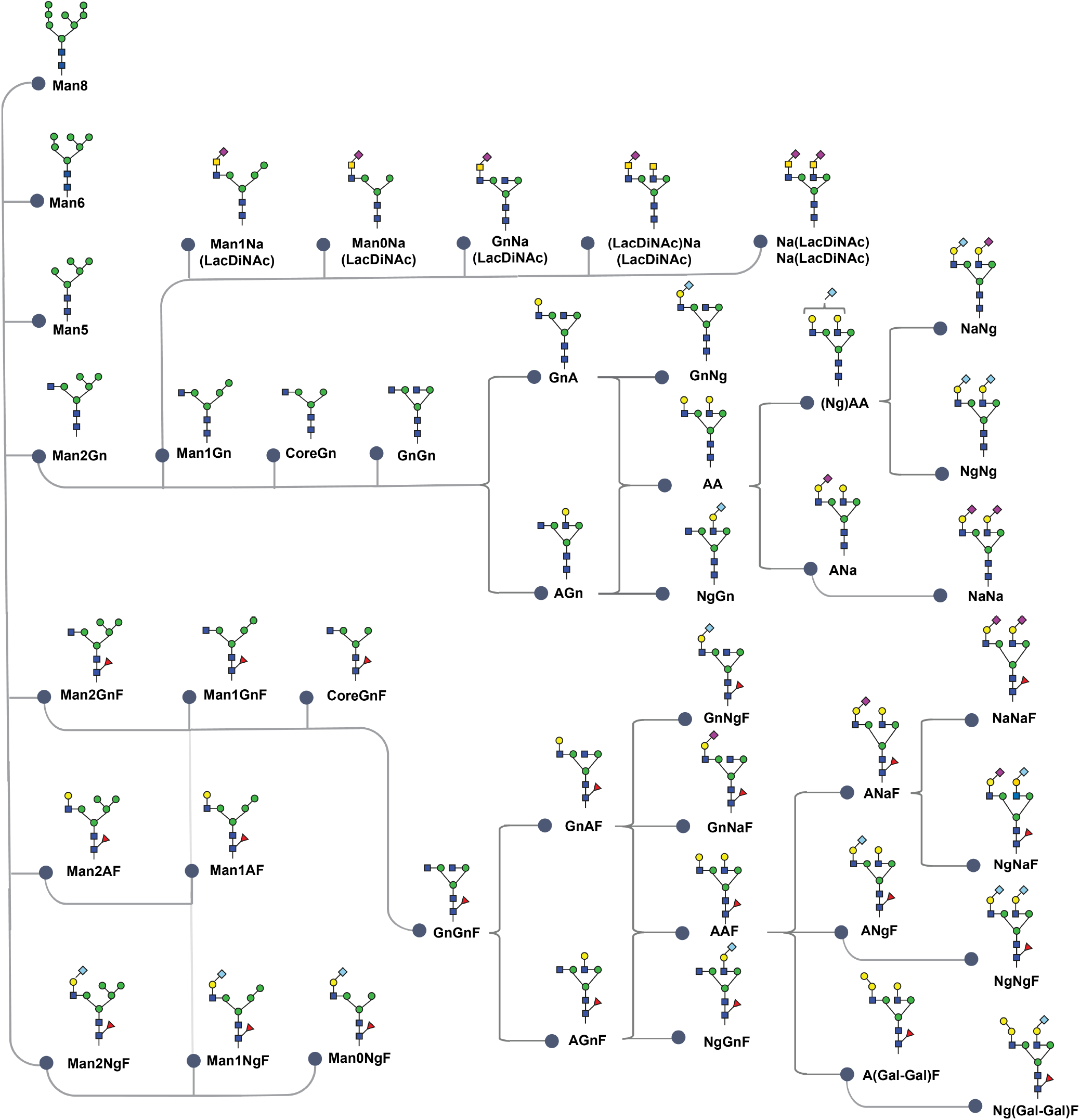
Consolidated representation of various sequential steps involved in the IgG N-glycosylation pathway. For simplicity, N-glycans carrying bisecting GlcNAc and α2-3 sialic acid linkage were omitted. The initial steps of N-glycosylation occurring within ER are highly conserved, but further downstream processing occurring in the Golgi exhibits a high species-specificity resulting in the synthesis of different (terminal) glyco-epitopes. In mammals, over 250 glycosyltransferases are known to reside within the Golgi ^40^. Their mission is to catalyze the transfer of a specific activated monosaccharide on a particular non-reducing end monosaccharide that is part of a larger glycan molecule. The specificity of individual glycosyltransferases is not only defined by the respective monosaccharide that is being transferred but also by the specific glycosidic linkage this monosaccharide is attached to. PGC-nLC-ESI-MS/MS analysis of N-glycans released from in vivo produced glycoproteins allows a highly representative interrogation of glycosidic pathway reactions as they occur within an organism that can be much more informative compared to the use of defined, synthetic glycan standards that might not ideally reflect the in vivo substrate. The ability to precisely differentiate these glyco-epitope features as such linkage, position, type of monosaccharide within a single analytical experiment provides a unique opportunity to interrogate the N-glycosylation pathways across different species.

Glyco-epitopes such as fucose (as in blood group antigens such as A, B, H, Lewis a, etc.) and/or sialic acids (Na, Ng) are known to be involved in host-pathogen interactions^38^. Our results clearly demonstrate that our GlycoRRT-PGC-LC-MS/MS strategy can be successfully applied to study the glyco-space in less well characterized species to understand the role of glycans in interspecies transmission of pathogens. As a first step, the IgG glyco-epitope map across 13 species presented here will augment our understanding of the molecular mechanisms of cross-species transmittable diseases, where species-specific glycosylation features play undoubtedly an important role ^39^.

The glycan spectral matching and scoring tool (GlycoRRT) including both, negative mode fragmentation and relative retention time, represents an important step forward towards reliable automated glycan identification. Overall, the library now contains more than 200 annotated N*-*glycans structures derived from synthetic N*-*glycan standards and human Immunoglobulins, plasma, blood, and urine. All N-glycan structures present in the library were characterized using (i) molecular monoisotopic mass and (ii) CID-MS/MS *de novo* sequencing. The results demonstrated the general applicability of the PGC-nLC-ESI MS/MS approach for detailed qualitative and quantitative glycomics analyses across species.

## Supporting information

Supplementary Results

## ACKNOWLEDGMENTS

We thank Max Planck Society for their generous financial support. KA was supported by PhD scholarship from Beilstein-Institut. This work was supported in part by Griffith University, European Union Seventh Framework Programme *IBD-BIOM* (project grant number 305479) and *Glycoproteomics* (project grant number 293847) and Australian Research Council Future Fellowship FT160100344 (to DK).

