## Supplementary Results for "Better together – relative retention time plus spectral matching improves automated glycan characterization using PGC-nLC-IT-ESI-MS/MS"

***Corresponding Authors**

**ORCID**

| | | |
|---|---|---|
| KA | – | 0000-0002-7596-5558 |
| AD | – | 0000-0002-3795-0291 |
| FA | – | 0000-0002-0112-7877 |
| PHS | – | 0000-0003-3394-8466 |
| MVI | – | 0000-0001-6302-7524 |
| NHP | – | 0000-0002-7532-4021 |
| DK | – | 0000-0002-8452-1350 |

#### ABSTRACT

Porous Graphitized Carbon nano-liquid chromatography tandem mass spectrometry (PGC-nLC-MS/MS) is a glycomics technique with the unique capacity to differentiate isobaric glycans. The lack of suitable software tools integrating chromatography and MS-information delivered by PGC-nLC-MS/MS has been limiting fast and robust glycan identification and quantitation. We report a LC-system-independent strategy called GlycoRRT that combines relative retention time (RRT) and negative ion fragment spectra analyses for isobaric structure-specific glycomics of PGC-nLC-MS/MS data. The GlycoRRT toolset is fully customizable and easily adaptable enabling semi-automated high-throughput structural assignments. The current library contains over 200 entries and their individual meta-data (MS instrumentation, experimental conditions, retention times, fragmentation profiles and glycan structural diagnostic ion features) relevant for reliable data analyses. The GlycoRRT workflow was employed to map the N- and O-glycome in blood group matched human plasma and urine as well as decipher Immunoglobulin (IgG) glycosylation features from 13 different animal species. We have also developed visualization tools to enable a consistent, reliable, and reproducible analysis of large sets of multidimensional PGC-nLC-MS/MS glycomics data. This comprehensive glycan resource provides the glycan map of human and animal species, will serve as a reference in dissecting the role of glycans in host pathogen interaction and zoonotic disease transmission.

**Key words:** Glycomics - Porous Graphitized Carbon – Immunoglobulin G – Mass spectrometry – Plasma glycomics

#### *ONLINE MATERIALS AND METHODS*

Blood group matched plasma and urine, animal plasma, pooled whole blood and plasma were obtained from Athens Research & Technology (Athens, GA, USA). Human and animal IgG's was purified from plasma samples by Melon Gel purification according to the manufacturer's protocol. Briefly, serum samples were diluted 1:10 and the diluted serum was added to a minispin column containing the Melon Gel resin. After 5 min incubation, the purified IgG was collected in the flow through following the manufacturer's instructions.

##### *SDS-PAGE and electro-blotting*

Five µg of protein (Human Immunoglobulins and IgG isolated from the animal serum) were incubated with 4 µL 4x SDS-PAGE sample buffer (0.25 M Tris/HCl pH 6.8, 40% [v/v] Glycerin, 4% [w/v] SDS, 0.015% (w/v) Bromophenol blue) containing 50 mM dithiotreithol (DTT) at 96°C for 5 minutes. After cooling down to room temperature, an aliquot of iodoacetamide solution (500 mM) was added to a final concentration of 50 mM iodoacetamide and incubated for 30 minutes in the dark before the proteins were subjected to SDS-PAGE separation on a precast 10% Mini-PROTEAN® TGX™ Gel (Biorad). Electrophoresis was performed at 200 V until the indicator band reached to bottom end of the gel. After SDS-PAGE, proteins were subsequently semidry electro-blotted onto PVDF membranes (0.2 µm pore size, Biorad, Munich, Germany) using the Trans-Blot® Turbo™ Transfer System (Biorad). Blotting was performed according to manufacturer's recommendations (7 min at 2.5 A and 25 V). Transferred proteins were visualised after staining with direct blue 71 as described previously (1).

##### *Dotblot*

A methanol-soaked tissue (e.g. Kim-Wipe) was placed into a glass petri dish. A piece of Immobilon-P Transfer PVDF Membrane was placed on top of the tissue and was wetted with MeOH. The upper surface of the PVDF membrane was allowed to dry. From the protein sample solution 3 µL was spotted in 1 µL volumes onto the PVDF membrane. After each step the droplet was allowed to dry. The tissue und the membrane was kept wet until all spots were applied. The membrane was allowed to dry over night

##### *Glycan release*

Direct blue 71 stained bands were cut from the PVDF membrane and transferred into 96 well plates for sequential N- and O-glycan release from the PVDF membrane as described previously (1). Any free PVDF surface was blocked with polyvinylpyrrolidone solution (1% PVP 40 in 50% methanol) and incubated over night at 37°C using 15 µL N-Glycosidase F (PNGase F, Roche Diagnostics, Mannheim, Germany) solution (0.17 U/µL in 10 mM $NH_4HCO_3$). *N*-glycans were collected and reduced for 3 hours in 1 M $NaBH_4$ in 50 mM KOH.

The N-glycan free sample on the PVDF membrane was subjected to a chemical *O*-glycan release via reductive beta-elimination (0.5 M $NaBH_4$ in 50 mM KOH, 50°C for 16 hours) and collected from the 96-well plates as described previously (1, 2). All samples were subsequently desalted by cation exchange chromatography (Dowex 50wX8, Biorad) using self-made micro spin columns as described in detail previously. After desalting, samples were further subjected to a carbon clean up via PGC micro-spin columns.

***PGC clean up***

A filter TopTip (Glygen, Columbia, MD) was filled with PGC material (resin was obtained from an Alltech Extract-Clean™ Carbograph SPE Column, Deerfield, IL). The micro spin column was washed three times with 50 µL 80% acetonitrile containing 0.1% TFA and subsequently equilibrated by washing three times with 50µL 0.1% TFA. The samples were loaded and the columns washed two times with 50 µL 0.1% TFA before glycans were eluted using 2x50 µL 80% acetonitrile containing 0.1% TFA. Eluted glycan samples were dried in the Speed Vac concentrator without any additional heat. Glycan samples were then reconstituted in 20 µL water for PGC nano liquid chromatography - ESI MS/MS analysis, a 3 µL aliquot was injected.

***PGC NANO LC-ESI IT-MS/MS ANALYSIS***

PGC-LC was performed on an Ultimate 3000 UHPLC system (Dionex, Part of Thermo Fisher, Germany) online coupled to an amaZon speed ETD ion trap mass spectrometer (Bruker Daltonics, Bremen, Germany). The LC system was connected to the mass spectrometer using the nano-flow ESI sprayer (Bruker Daltonics, Bremen, Germany). The instrument was controlled using Hystar 3.2 software.  The spectra were analysed using Compass Data Analysis v4.2 (Bruker Daltonics, Bremen). The instrument was set up to perform CID fragmentation on the selected precursors. An *m/z* range from 350-1600 Da was used for data dependant precursor scanning. The three most intense signals in every MS scan were selected for MS/MS experiments. MS as well as MS/MS data were recorded in the instrument's "Ultra scan mode" and "Enhanced resolution mode" respectively. Specific instrumental operational parameters used in the present investigation are listed in Table 2.2. Glycans were loaded onto a PGC (porous graphitized carbon) precolumn (Hypercarb KAPPA 30 x 0.32 mm, 5 µm particle size) and separated on an analytical PGC column (Hypercarb™ PGC Column, 100 mm x 75 µm particle size 3 µm, both ThermoFisher Scientific, Waltham, MA). The samples were loaded onto the precolumn at a flow rate of 6 µL/min in 98% buffer A (10 mM ammonium bicarbonate). The starting conditions for the analytical column at a flow rate of 1 µL/min were 3% buffer B (10 mM ammonium bicarbonate in 60% acetonitrile). The gradient conditions were as follows: increase of buffer B from 3 to 16% (5.5-7.5 min), further increase to 40.0% B (7.5-

55.5 min), followed by a steeper increase to 90% B (55.5-62.0 min). The column was held at 95% B for 8 min (62-70 min). At the same time the precolumn was flushed with 90% Buffer C (10 mM ammonium bicarbonate in 90% acetonitrile) at a flow rate of 6 µL/min before reequilibrating the precolumn as well as the analytical column in 98% buffer A for 7 minutes. All analyses were performed in triplicate.

##### *SPECTRAL LIBRARY DESIGN AND IMPLEMENTATION*

The library was created using LibraryEditor V4.2, which is a part of Compass Data Analysis v4.2 (Bruker Daltonics, Bremen) and Excel (Microsoft). The library features meta data features such as absolute chromatographic retention time of the reduced *N*-glycan alditols, mass (*m/z*) and tandem MS spectra.

###### *LC-MS Data*

The extracted ion chromatogram (EIC) for each glycan was derived based on the glycan $[M–H]^{-}$/$[M–2H]^{2-}$ masses. Absolute retention times of the EIC of the given *m/z* values under the experimental conditions were recorded and exported to the library.

###### *MSMS Spectral Library*

To simplify the product ion spectra obtained for each standard *N*-glycan, peaks in each MSMS spectra were merged with a mass tolerance for the precursor (0.3 Da). The observed product ions over the threshold 25% were exported to LibraryEditor spectral library including the information about the observed diagnostic fragment ions, theoretical glycan mass and glycan composition represented in the GlycoMod format. Also, each entry also contained a glycan name represented in Proglycan nomenclature to distinguish the structural isomers in a single letter code sequence.

###### *Library Evaluation by Spectral Searching*

Before implementing the GlycanRRT library in the glycan analysis workflow, the datasets used to create the library was re-searched the GlycanRRT library. Relative Retention time (RRT) was calculated as the ratio of absolute retention time of the glycan to the absolute retention time of the query using the expression RRT = (Sample RT / Standard RT)

###### *LC-MS/MS blind trial*

Three different LC-MS/MS glycan datasets from different participants were selected for the blind trial. The blind trial spectra were processed and then searched against all the spectra present the GlycanRRT library. The results were exported into excel containing all the meta data including % fit score.

##### *GLYCAN IDENTIFICATION AND RELATIVE QUANTITATION*

Glycans were automatically identified using an in house established GlycoRRT spectral database and the spectral library tool integrated in Compass Data Analysis 4.2 (Bruker). *N*-glycan structures not present in the database were manually annotated using Data Analysis 4.2. After identification of the *N*-glycans, relative quantitation was performed using the QuantAnalysis tool (Bruker), which determines the area under the curve obtained from the individual extracted ion chromatograms (EIC) from multiple analyses. The integration of every extracted ion chromatogram was validated manually, in particular for peaks with very low abundance. The values out of three technical replicates were averaged for this study and their standard deviations are represented in the error bars. In each sample the total amount of the identified glycan structures were taken as 100% and their relative distribution determined.

### SUPLEMENTARY RESULTS

#### *NEGATIVE ION MODE PGC-LC-ESI-MS/MS GLYCOMICS*

Fragment ions generated in the negative-ion MS/MS spectra are rich in information that assists meticulous glycan structural assignment. In contrast to positive ion fragmentation, in negative ion fragmentation diagnostic ions are produced by a single pathway following proton abstraction from the specific hydroxyl groups (3). Thus, the observed diagnostic fragment ions derived from negative ion MS/MS spectra can be used to elucidate structural features such as the position of fucose, the core type of O-glycans and the branching of N-glycan structures (4-6). The most important diagnostic ions observed upon CID of negatively charged molecules are highly informative D-ions (providing information on the 6 arm antennae substitutions) and E-ions (3 arm antennae). The presence or absence of a bisecting GlcNAc can be inferred from the D-221 ions. C-ions derived from the non-reducing end of the glycan provide information on the sequence of the constituent monosaccharide residues (**Figure S1**). The presence of an abundant [D-221] ion (e.g., *m/z* = 508 or 670 or 961), for example, indicates the presence of a bisecting GlcNAc residue whereas as the Z1 (*m/z* = 350) and Z2 (*m/z* = 553) diagnostic ions are indicative for the presence of core fucosylation (**Table S3, Figure S2**).

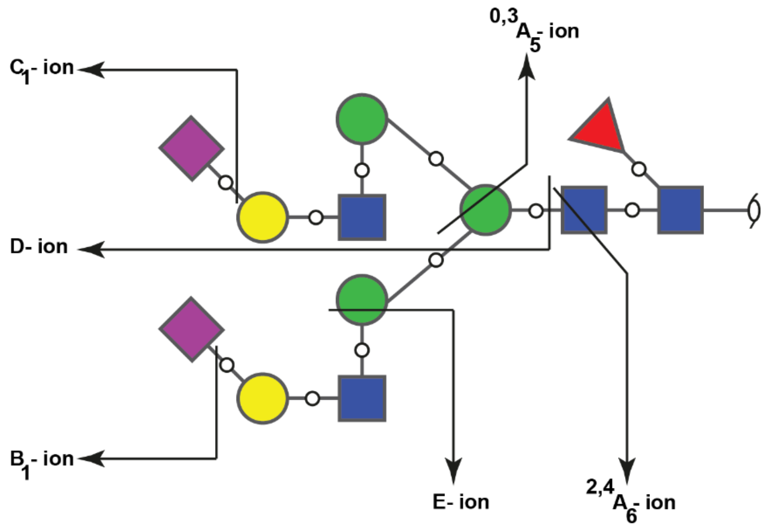


***Figure S1: Exemplary bi-antennary N-glycan showing the major fragment ions.*** *The oxygen atoms involved in the glycosidic bond are shown to differentiate B- and C-ions.*

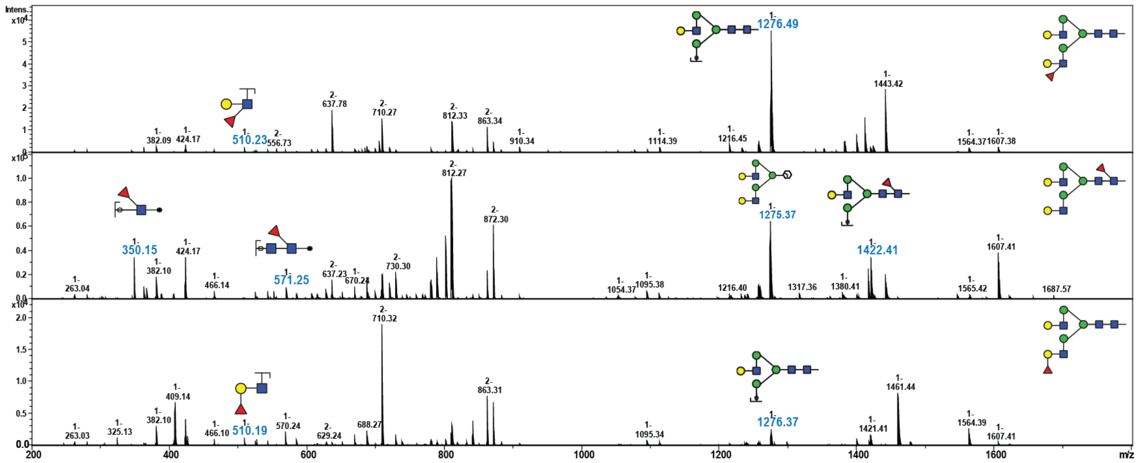


***Figure S2:*** *Characterisation of neutral fucosylated* N-*glycan structural features by negative ion PGC-nLC-ESI-IT-MS/MS. The presence of core fucose is indicated by Z1 ion at* m/z *350 and Z2 ion at 571, whereas the presence of arm fucosylation can be differentiated by B and C ions at* m/z *510 and 528.*

#### *GLYCORRT SPECTRAL LIBRARY*

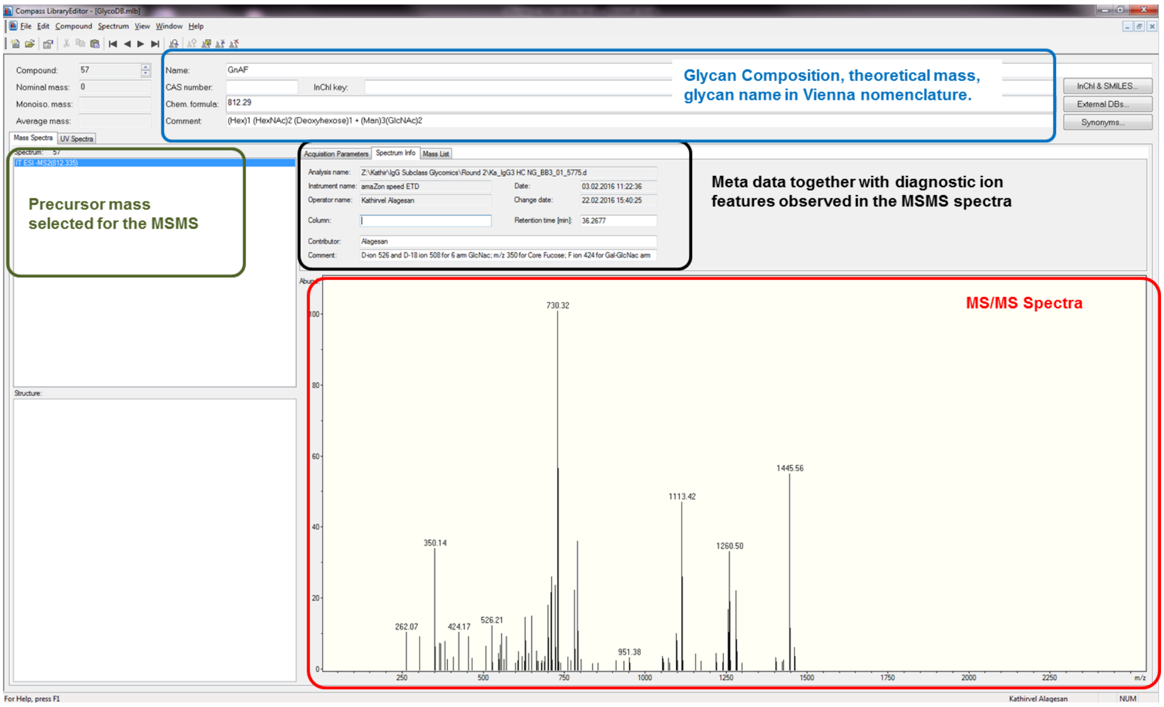


***Figure S3:*** *GlycoRRT MS library screenshot example for the mono galactosylated fucosylated* N-*glycan (GnAF). The GlycoRRT library developed contained data derived from 33 synthetic N-glycan standards (**Table S1**) that included the absolute retention time value for each individual glycan together with the respective negative mode product ion spectra and the observed diagnostic fragment ions, theoretical glycan mass and glycan composition represented in the GlycoMod format.*

##### ***Spectral matching automation in data analysis***

The identification of previously observed glycans was accelerated by automated spectral matching. The Bruker software LibraryEditor was connected with DataAnalysis and was used to build the libraries from the recorded fragmentation data. The library was then used to identify glycans in new sample files using the Identify – Spectra command in DataAnalysis. To facilitate, processing of large samples, a data analysis method was created to enable automated compound creation and identification (***See Script below***)

```
'This script contains a typical LC/MS/MS processing

option explicit

Analysis.compounds.clear

dim filename, Filename2, targetxml, targetmgf

'select a retention time range of 10 - 65 min

Analysis.AddChromatogramRangeSelection 10, 65, 0, 0

'perform Find Compounds AutoMS(n)

Analysis.FindAutoMSn

'deconvolute, search and also export to MGF

Analysis.Compounds.Identify

form.close
```

***Glycan elution Rules of PGC***

Specific glycan structure features are known to influence the PGC elution behaviour. The order of elution of individual structural isomers is also highly conserved (7). N-glycans carrying a bisecting *N*-acetylglucosamine (GlcNAc) elute earlier than their non-bisected structural isomers**.** Similarly, presence of core fucosylation increases retention whereas Lewis type antenna-fucosylation results in shorter retention times (**Table S6**). The polar retention effect accounts for increased selectivity, in particularly for sialylated glycans. Very subtle changes such as the linkage position of *N*-acetylneuraminic acid residues significantly alter the retention time of isobaric glycan structures. The glycan structures carrying α2,3-linked NeuAc residues are retained much stronger compared to their α2,6-linked counterparts. In addition to their different elution time, the in negative mode MS/MS spectra of the PGC separated compounds also provide additional confirmation on the NeuAc linkage positions (**Figure S4**). The high separation power of PGC-LC is just not limited to sialylated residues but also equally applies to other *N*-glycan classes such as high mannose isomers and complex *N*-glycans structures carrying Lewis x/a/y/b structures. Thus, combining the high separation capacity offered by PGC with information rich negative mode fragmentation represents a unique technology that acquires multidimensional data that all together provides profound structural and quantitative information also from very complex sample sources while using a minimum of initial sample – a crucial prerequisite for any glycan sequencing technology that is to be applicable for clinical research.

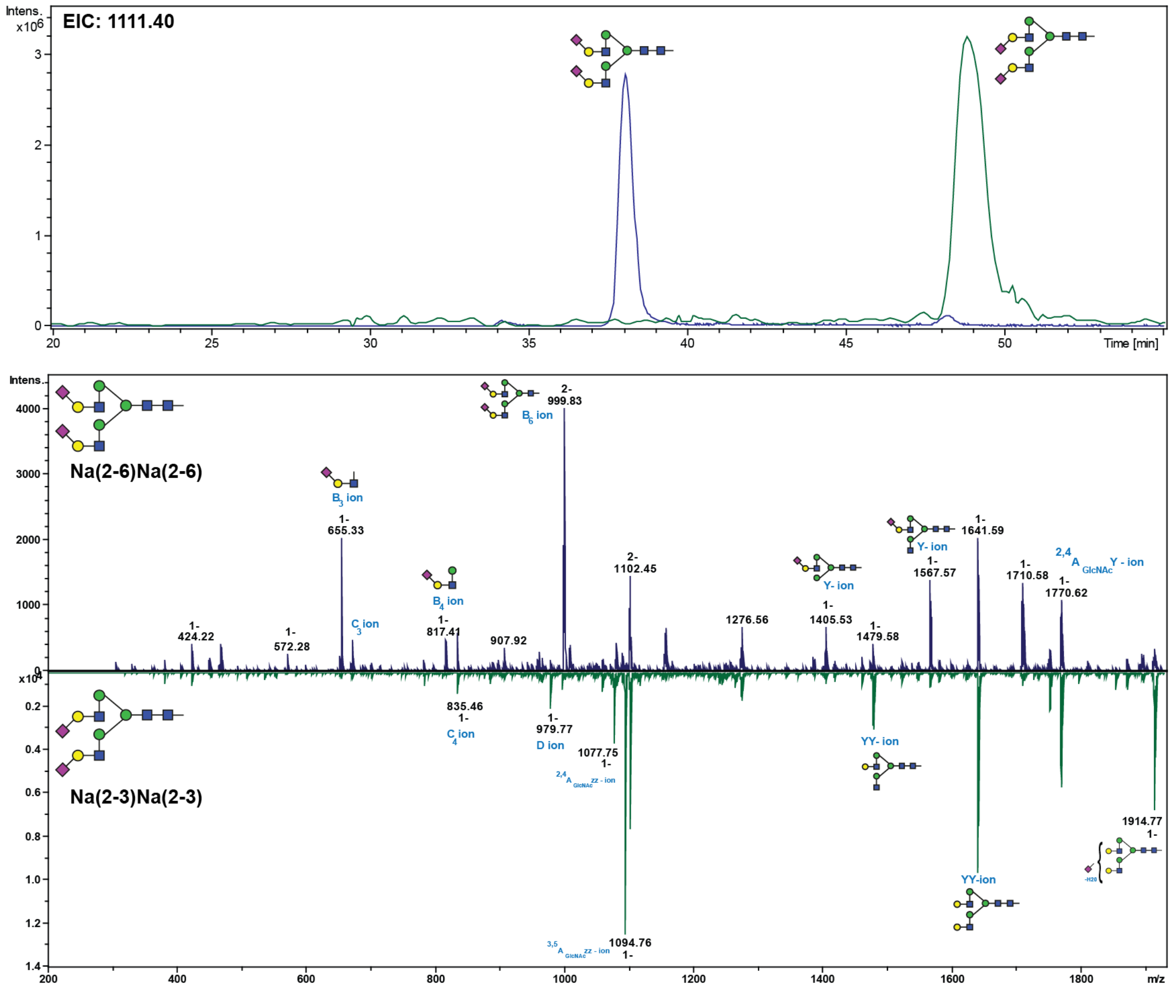


***Figure S4:*** *PGC-LC separation of α2,6 and α2,3 linked Neu5Ac isomers. Extracted ion chromatogram (EIC) of $[M-2H]^{2-}$ = 1111.40 corresponding to the composition $Hex_2HexNAc_2NeuAc_2$ + $Man_3GlcNAc_2$ eluting at two different time points (Top panel). In addition to different elution profiles, the respective negative mode MS/MS spectra showed structure specific MS/MS signatures that allowed isomer differentiation. The prominent water loss (-18 Da) observed for the α2,3 linked Neu5Ac* (m/z *1914.77) indicated the presence of a α2,3 linked Neu5Ac (bottom panel).*

***Factors influencing glycan separation by PGC-LC***

Although the exact molecular nature of the retention mechanism leading to isobaric glycan separation by PGC-LC is yet still not fully understood, hydrophobic, polar and ionic interactions have been identified to contribute to analyte retention (8, 9). Unlike hydrophilic-interaction liquid chromatography (HILIC) and ion pairing reversed-phase chromatography (IP-RPC), glycan retention in PGC cannot be correlated to the monosaccharide compositions (hydrophilicity of the glycan) by multiple linear regression analysis (10). On the other hand, factors such as electrosorption, solvent, temperature and ion polarity are also known to influence the separation behaviour of sialylated glycans (11). Yet, little is known about the influence these parameters have on the isomer separation capacity offered by PGC. Thus the influence the column temperature has on the isomer separation capacity was evaluated using AGnF and GnAF *N*-glycans. In agreement with previous studies(11), higher column temperatures did result in an increased glycan retention capacity (**Figure S5**). Besides, the here presented data also indicates that an increase in column temperature also resulted in better baseline separation of the AGnF and GnAF isomers as evident from the *m/z* 812.33 extracted ion chromatogram (EIC) corresponding to doubly charged signal of these two standard glycans (**Figure S5**).

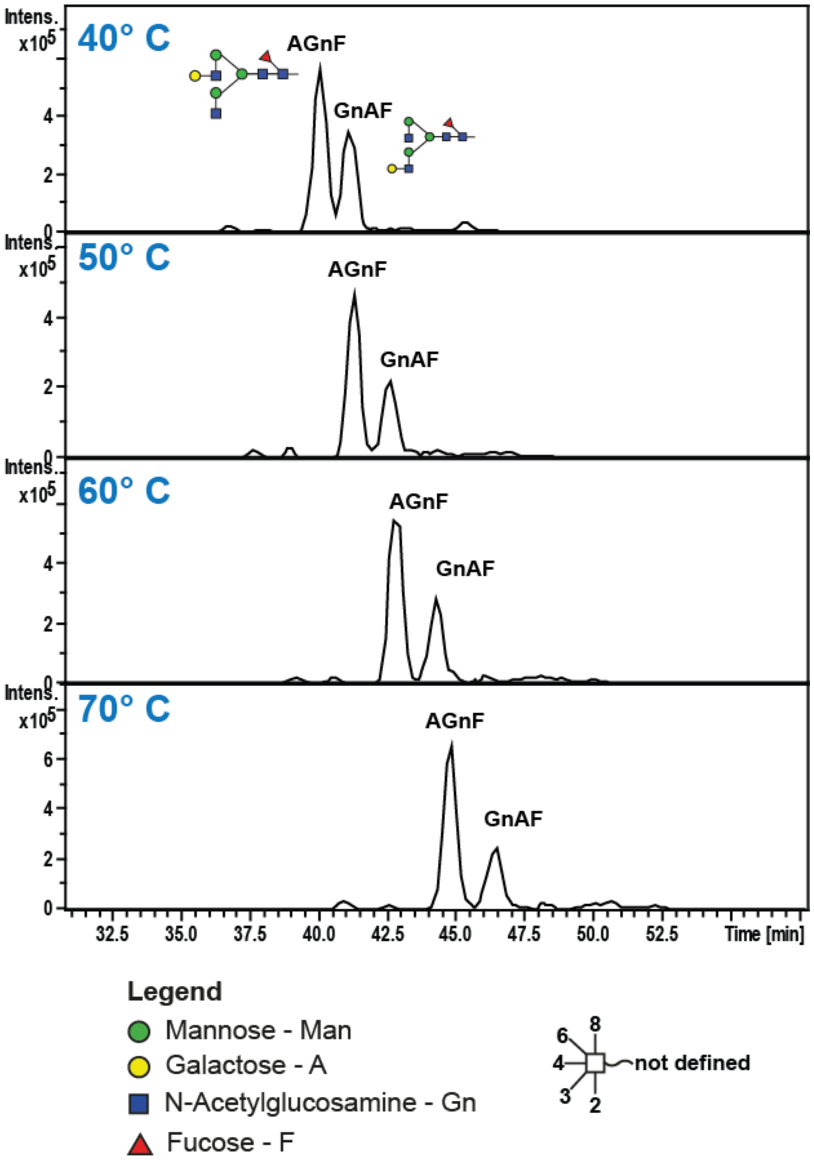


***Figure S5:*** *Influence of column temperature on isobaric glycan separation by PGC-LC. The retention of* N-*glycans clearly increased with higher column temperatures, which also resulted in better baseline separation of these two arm isomer* N-*glycan structures.*

In positive mode, applied electrospray voltage is known to cause polarisation of the carbon surface resulting in a total retention of sialylated glycans. This unwanted effect could be avoided by column grounding (11, 12). Therefore, the glycan separation behaviour of grounded and ungrounded columns in negative ionisation PGC-LC ESI MS/MS was evaluated using standard *N*-glycans derived from bovine fetuin. While the elution pattern of neutral and mono-sialylated glycans were not affected by the grounding status, tri- and tetra-antennary sialylated *N*-glycans were retained slightly stronger in non-grounded columns. On contrary to positive mode PGC-LC, where a complete retention of these highly sialylated *N*-glycans was reported. The present negative ion mode PGC-LC ESI MS/MS system used in the analysis did not result in any such effect (**Figure S6**).

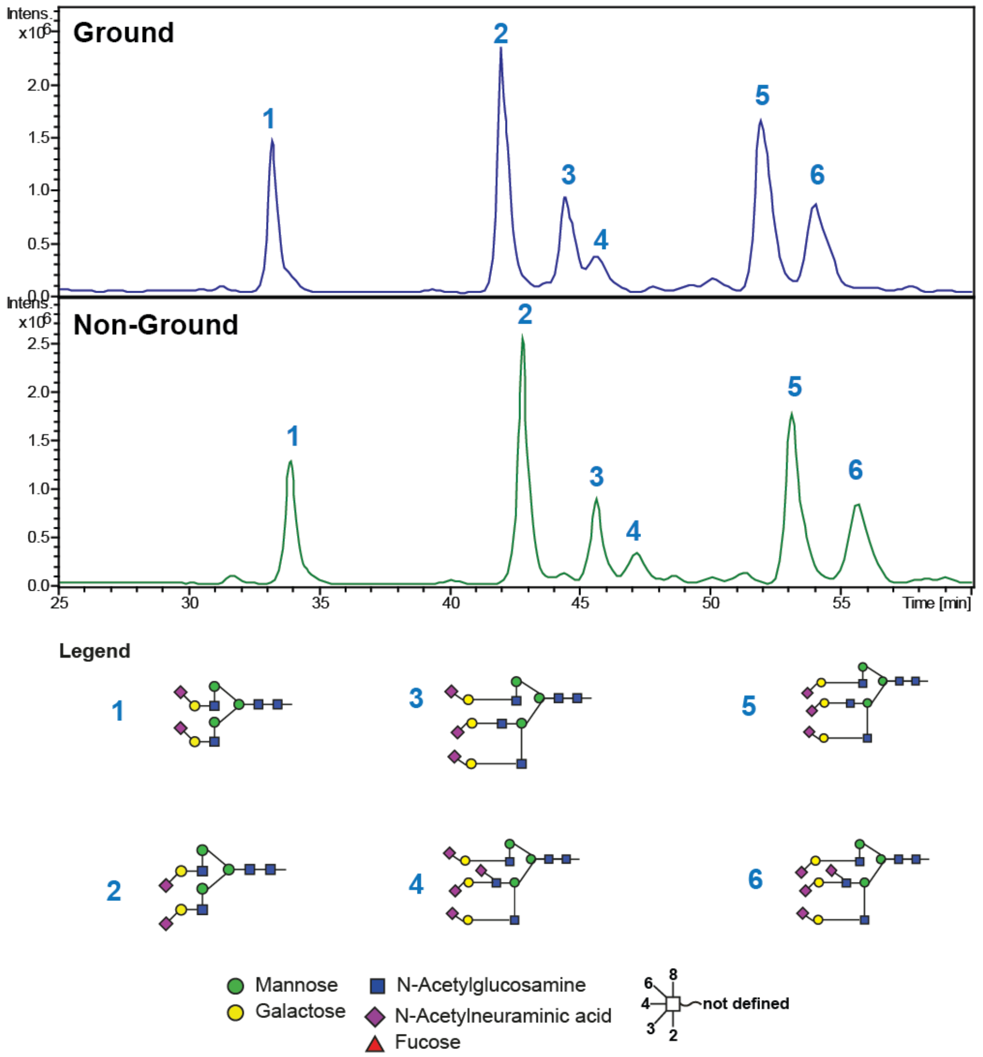


***Figure S6:*** *Influence of column grounding on* N*-glycan retention in PGC-LC. Sialylated* N*-glycan alditols are retained slightly stronger in non-grounded columns in comparison to grounded columns.*

These features (high column temperature and not grounded) could potentially be exploited for improving the analysis of weakly retained oligosaccharides. Very high temperatures, however, should be avoided in glycan analysis to reduce any potential analyte degradation (e.g. sialyl linkages) (8). Despite the absolute LC retention time shifts (either due to eluent or column batch differences); the PGC separation features to separate structural isomers remained highly reproducible**.** This unique retention behaviour in combination with individual signature fragment spectra recorded from negatively charged precursor ions was employed to develop a spectra library based matching tool for the automated identification of glycan structures – a first crucial step in the development of an automated glycan sequencing tool.

##### *Influence of SPS parameter on quantification*

The Smart Parameter Setting (SPS) available in the used ion trap instrument supports in the auto-adjustment of the acquisition mass window to a target *m/z* value of interest. Typically, the SPS target is set in the middle of the intended acquisition mass range. Next, we evaluated the impact of SPS target parameter on glycan identification and quantitation using the Man6 glycan via direct infusion MS analysis. The SPS target was sequentially increased in 100 Da steps from *m/z* 900 to 1400 and any in- or decrease in the signal intensities was expressed in relation to the signal intensity observed at SPS 900.

Under the selected MS parameter settings Man6 can be detected either as a singly or doubly charged precursor ion or in both the charge states. The acquired data provided clear evidence that the SPS target *m/z* has a direct influence on the detected charge state (**Figure S7-A & B**). This was also explained by the fact that alterations in SPS target did not only optimise the acquisition mass window but also the ion transfer. This became evident by the observed signal intensity decrease of the doubly charged precursor that was compensated by a signal intensity increase of singly charged precursor. The SPS target shift from *m/z* 900 to 1400 resulted in an approximately 5-fold decrease in the singly charged precursor signal intensity (**Figure S7-A)** whereas the doubly charged precursor signal intensity decreased 0.38-fold (**Figure S7-B)**. When both charge states were considered for quantitation, however, the overall relative signal intensity decreased by 0.73-fold when the SPS target *m/z* was increased from 900 to 1400 (**Figure S7-C)**.

Intrigued by this observation, a panel of available *N*-glycan standards was tested to evaluate the SPS parameter influence on signal intensity and subsequent relative quantitation. In the case of Man2 the shift in SPS target *m/z* from 900 to 1400 resulted in the gradual decrease in signal intensity (**Figure S7-D**). In the case of tetra-antennary *N*-glycan (GnGn)(GnGn) ($[M-2H]^{2-}$ = 861.32), however, the SPS target *m/z* increase from *m/z* 900 to 1100 already resulted in 1.67-fold increase in signal intensity. A further increase to 1200 resulted in a 1.73-fold increase. The subsequent increase in SPS target *m/z* to 1400 resulted in the gradual signal intensity decline that in the end just left a 1.26-fold increase (**Figure S7-E**). A similar trend was observed for the bi-antennary monosialylated *N*-glycan NaA ($[M-2H]^{2-}$ = 965.84) (**Figure S7-F**).

An increase in SPS target *m/z* from 900 to 1400 had a larger effect for the tetra-antennary, bisected *N*-glycan (GnGnGn)(GnGn)bi ($[M-2H]^{2-}$ = 1064.44) resulting in an approximate 6.8-fold signal intensity increase (**Figure S7-G**). Considering the observed exceptional SPS parameter dependant influence on signal intensities and detected precursor ion charge it is necessary to develop optimised analysis parameters that depend upon the sample type to avoid any under-representation of individual glycan structures. The acquired data indicated that the selection of a SPS target *m/z* oriented towards the higher end of the intended *m/z* scan range provided better quantitative results compared to the traditional mid-range SPS selection when the glycans over a large *m/z* range are to be covered.

The use of exoglycosidase enzymes in combination with PGC-LC-ESI-MS has been reported to improve the relative quantitation of tri and tetra-antennary *N*-glycan classes (13). However, this approach requires an array of exoglycosidase enzymes and substantial sample amount involving multiple sample preparation steps. Also, information pertaining to individual e.g. NeuAc structure isomers present in the undigested sample is lost. Considering the intrinsic complexity of the glycan, there is no single universal method currently available that will cover each and every aspect in glycan analysis. Thus, it is of utmost importance that a method provides suitable versatility to cover a wide *m/z* and structure range as good as possible. Depending on the target sample, conditions need to be optimised, but once determined for a specific sample set (e.g. body fluid, tissue) these optimised settings can be widely applied.

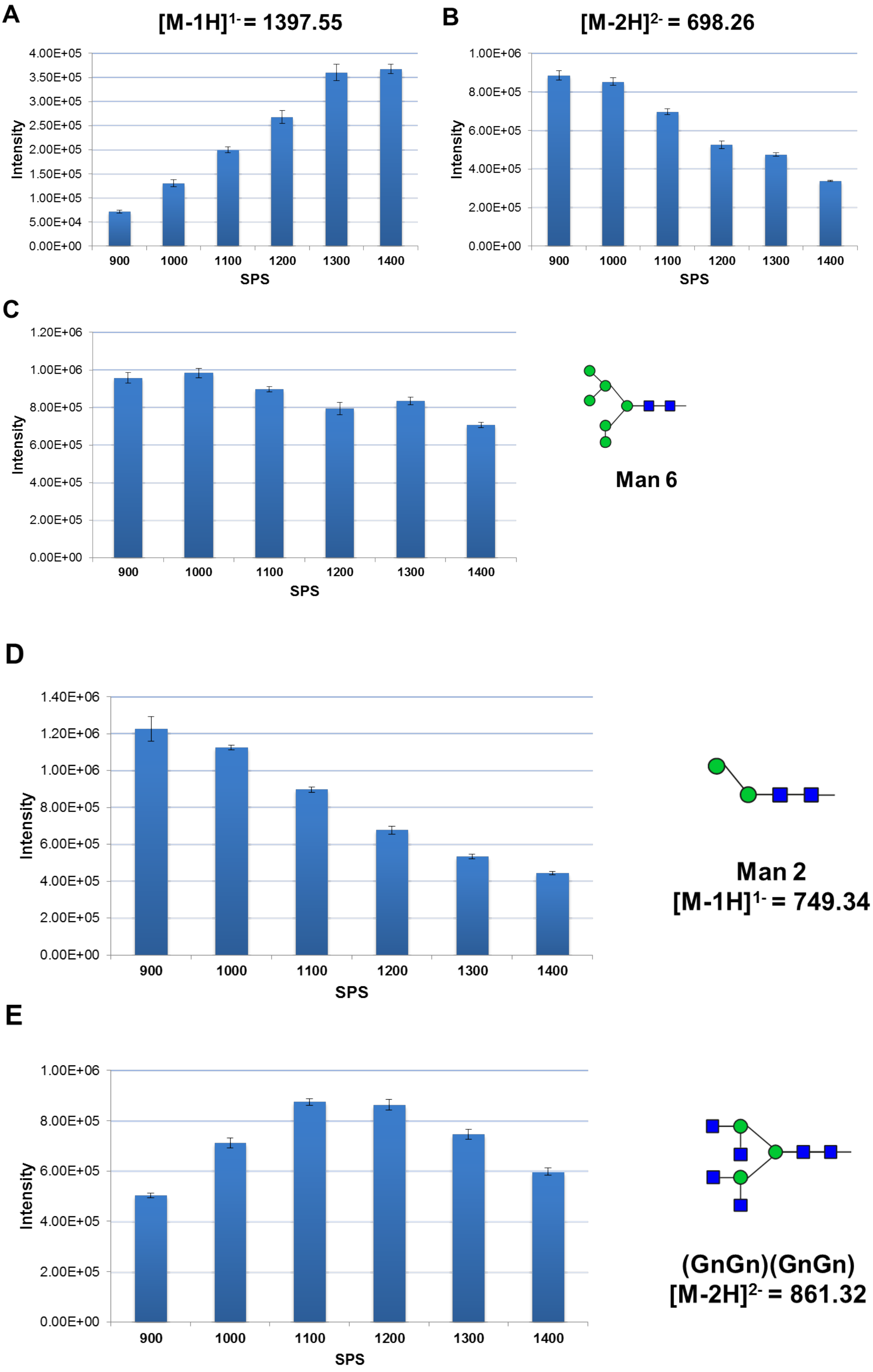
A
[M-1H]1- = 1397.55
Intensity
4.00E+05
3.50E+05
3.00E+05
2.50E+05
2.00E+05
1.50E+05
1.00E+05
5.00E+04
0.00E+00
900
1000
1100
1200
1300
1400
SPS
B
[M-2H]2- = 698.26
Intensity
1.00E+06
8.00E+05
6.00E+05
4.00E+05
2.00E+05
0.00E+00
900
1000
1100
1200
1300
1400
SPS
C
Intensity
1.20E+06
1.00E+06
8.00E+05
6.00E+05
4.00E+05
2.00E+05
0.00E+00
900
1000
1100
1200
1300
1400
SPS
Man 6
D
Intensity
1.40E+06
1.20E+06
1.00E+06
8.00E+05
6.00E+05
4.00E+05
2.00E+05
0.00E+00
900
1000
1100
1200
1300
1400
SPS
Man 2
[M-1H]1- = 749.34
E
Intensity
1.00E+06
8.00E+05
6.00E+05
4.00E+05
2.00E+05
0.00E+00
900
1000
1100
1200
1300
1400
SPS
(GnGn)(GnGn)
[M-2H]2- = 861.32

F

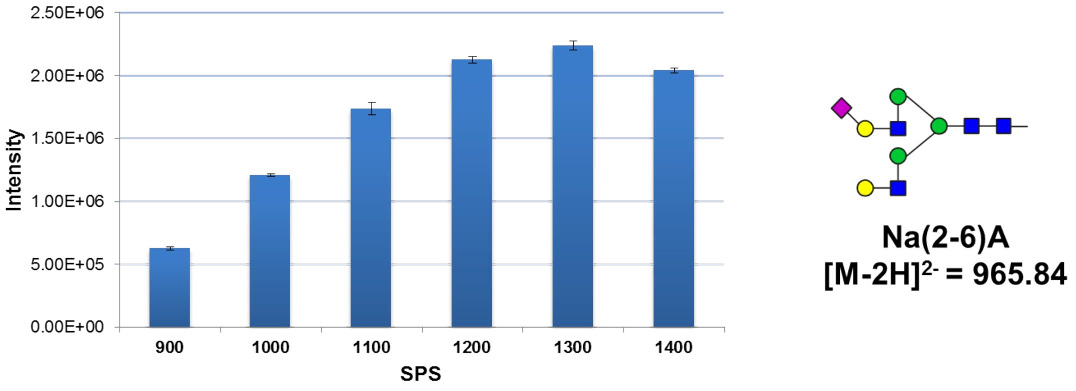


G

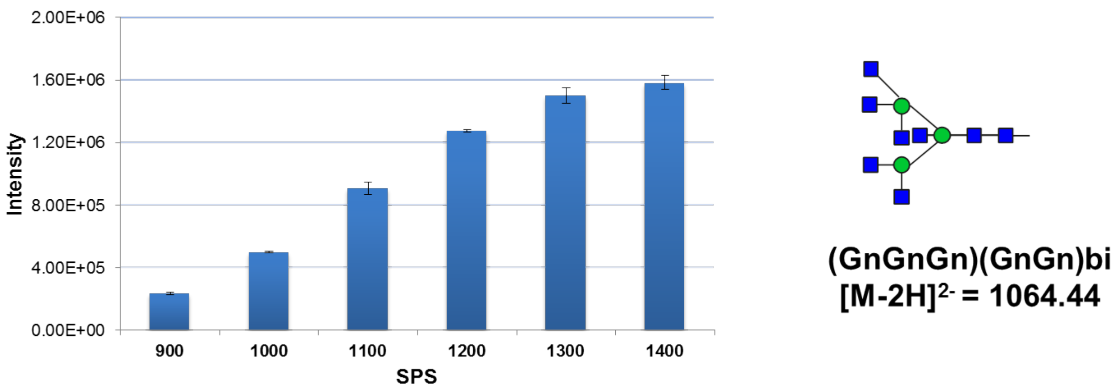


***Figure S7- A-G:*** *Influence of SPS target* m/z *on glycan quantitation evaluated by direct infusion MS analysis of reduced* N-*glycan alditols over the indicated mass ranges. The signal intensities of each* N-*glycan structure were denoted as the average isotope intensities summed up over 20 sec elution range. Mean and standard deviation of triplicate summed up intensities are shown.*

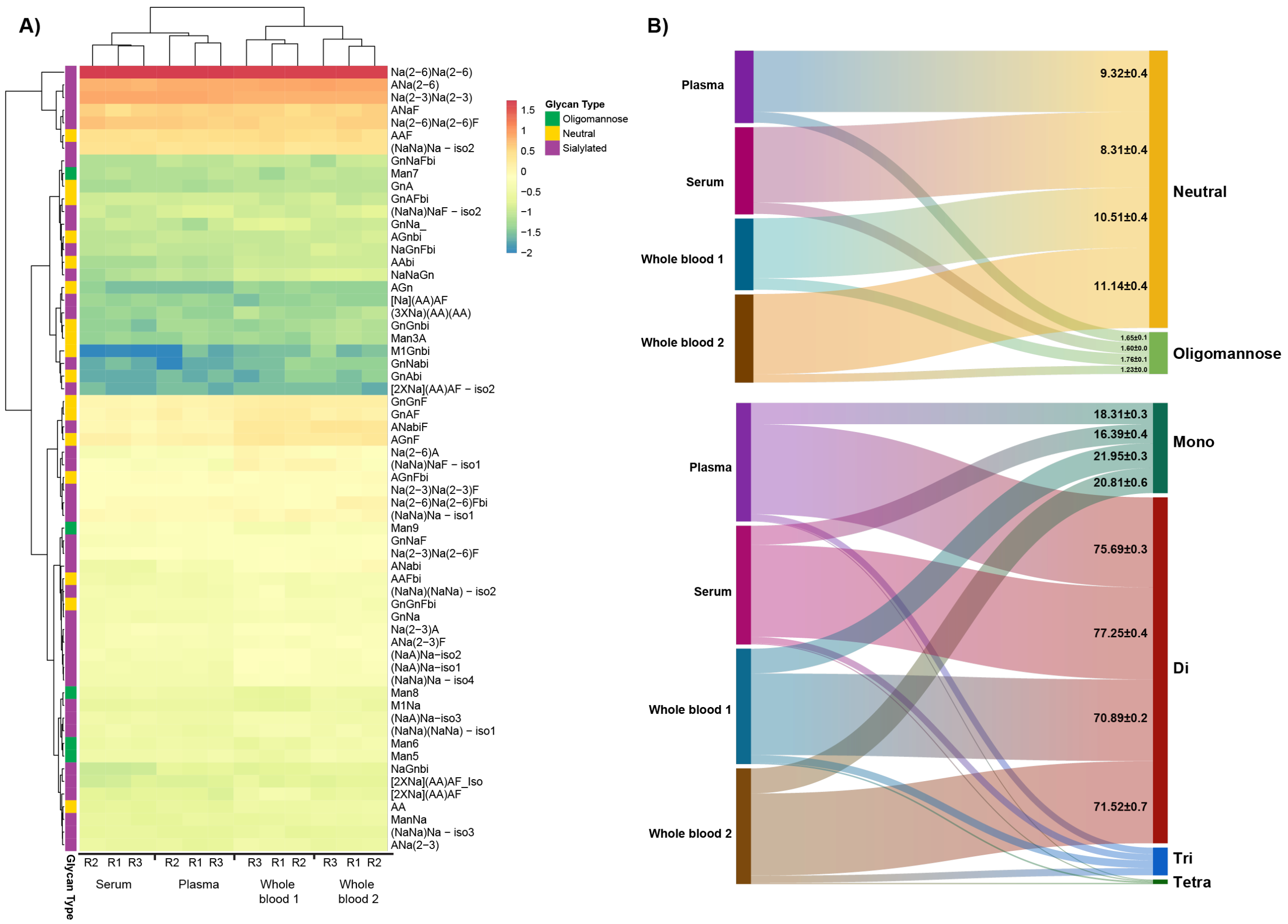


***Figure S8:*** *Comparison of* N*-glycans released from human serum, plasma and whole blood. Heat map of the log-transformed relative abundances of all detected* N*-glycans derived from human plasma, serum and whole blood. The samples were ordered by hierarchical clustering using the Euclidean distance matrix with complete linkage. Columns indicate glycan sample source and rows indicate the glycan structures (Supplementary table S7). Based on the glycan relative abundance data, the algorithm classified the samples into two distinctive classes containing: (i) serum & plasma and (ii) whole blood.*

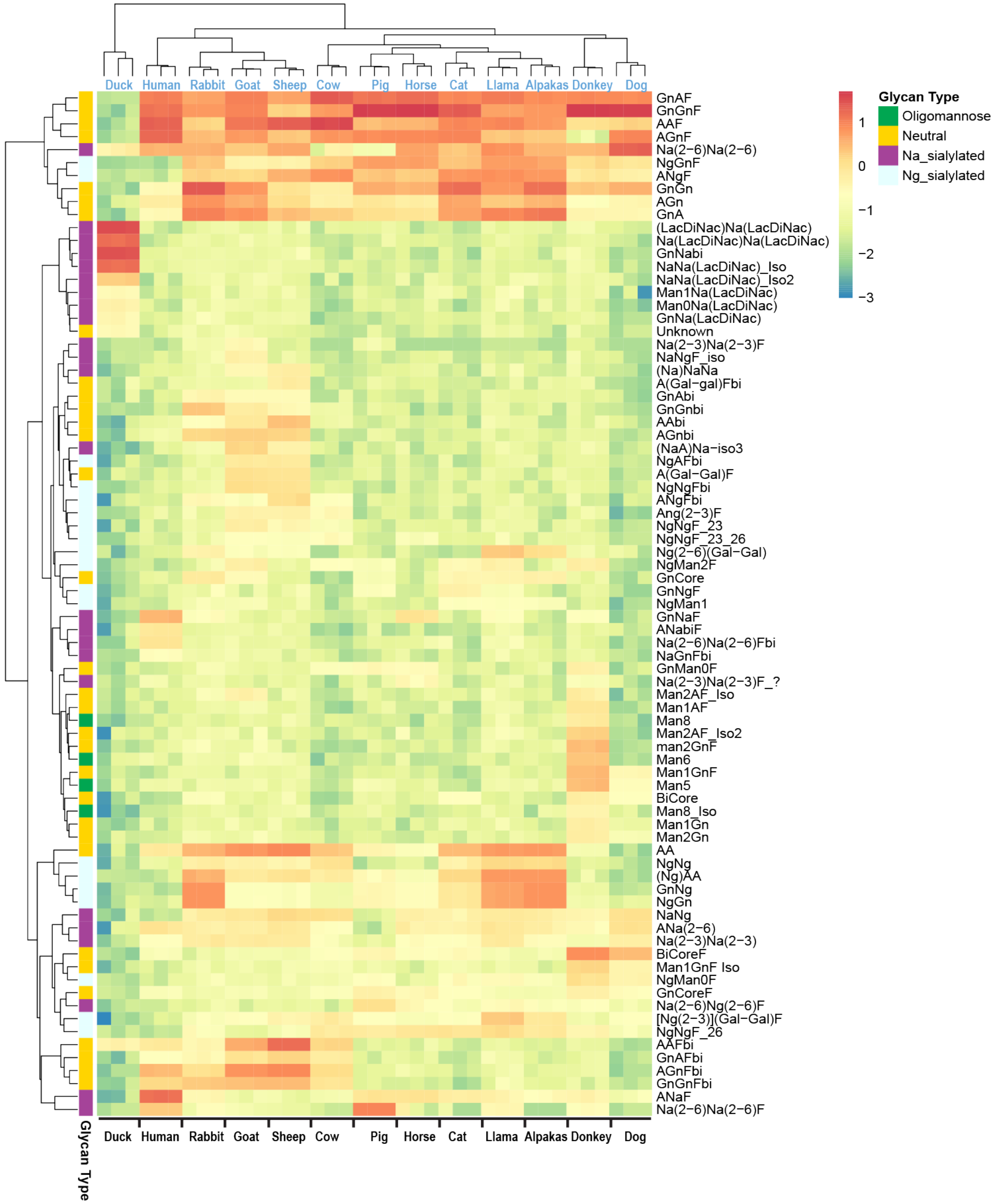


***Figure S9:*** *Heatmap of the logarithmic relative abundances of IgG N-glycans from twelve different mammals and one bird. Samples were ordered by unsupervised hierarchical clustering using the Euclidean distance matrix with complete linkage.*

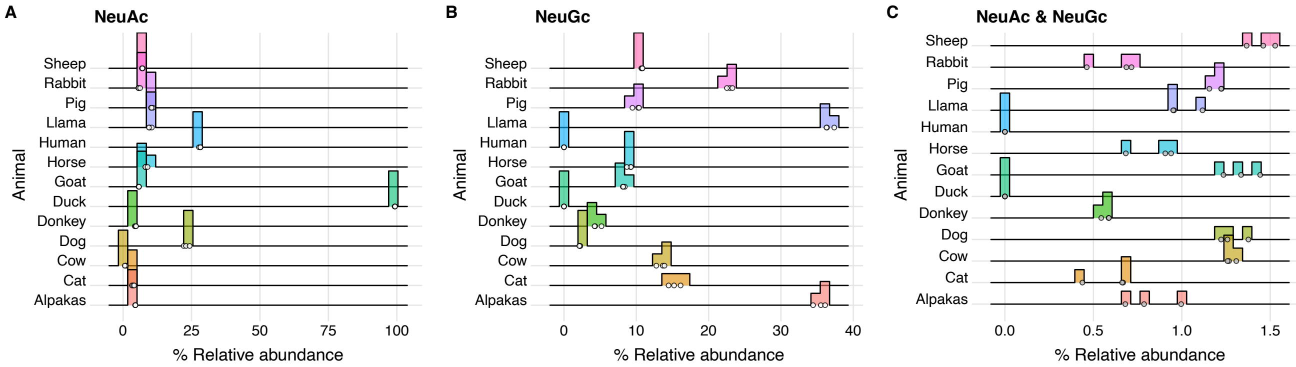


***Figure S10:*** *Ridgeline plot of relative abundances of IgG N-glycans containing sialic acids from twelve different mammals and one bird.*

##### *Data analysis and visualization pipeline*

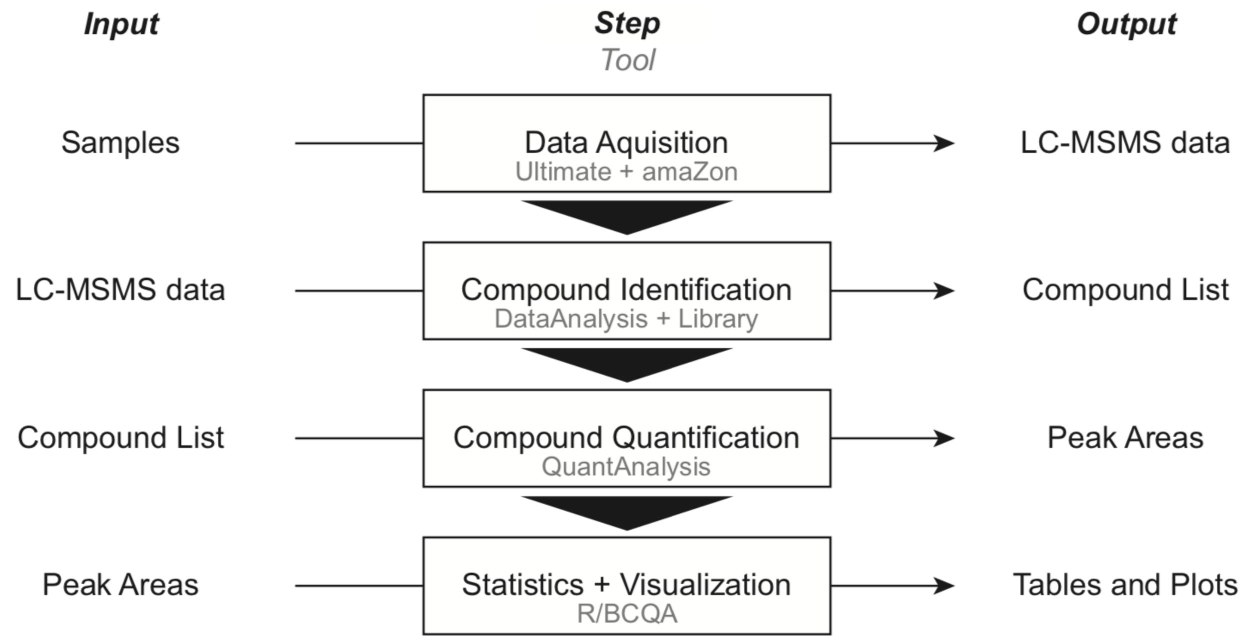


***Figure S11:*** *The glycomics data analysis was divided into four major procedures. The raw data was acquired on the LCMS instrument and glycan structures were identified manually in Bruker DataAnalysis from retention time, precursor mass and fragment spectra according to literature information. A spectra matching approach based on spectra libraries was established. The quantification was done automatically in Bruker QuantAnalysis and verified manually supported by custom-made software tools. The output of the quantification was normalized and evaluated using statistical methods according to the respective biological context of each sample set. Especially the last step was enhanced with own software (BCQA package) developed in the data analysis language and environment R. The results of the differential glycan expression is visualized as a heatmap, which describes the relationship of all significantly regulated glycans (rows) across all samples (columns). Hierarchical clustering is performed on both glycan (rows) and sample (columns) levels.*

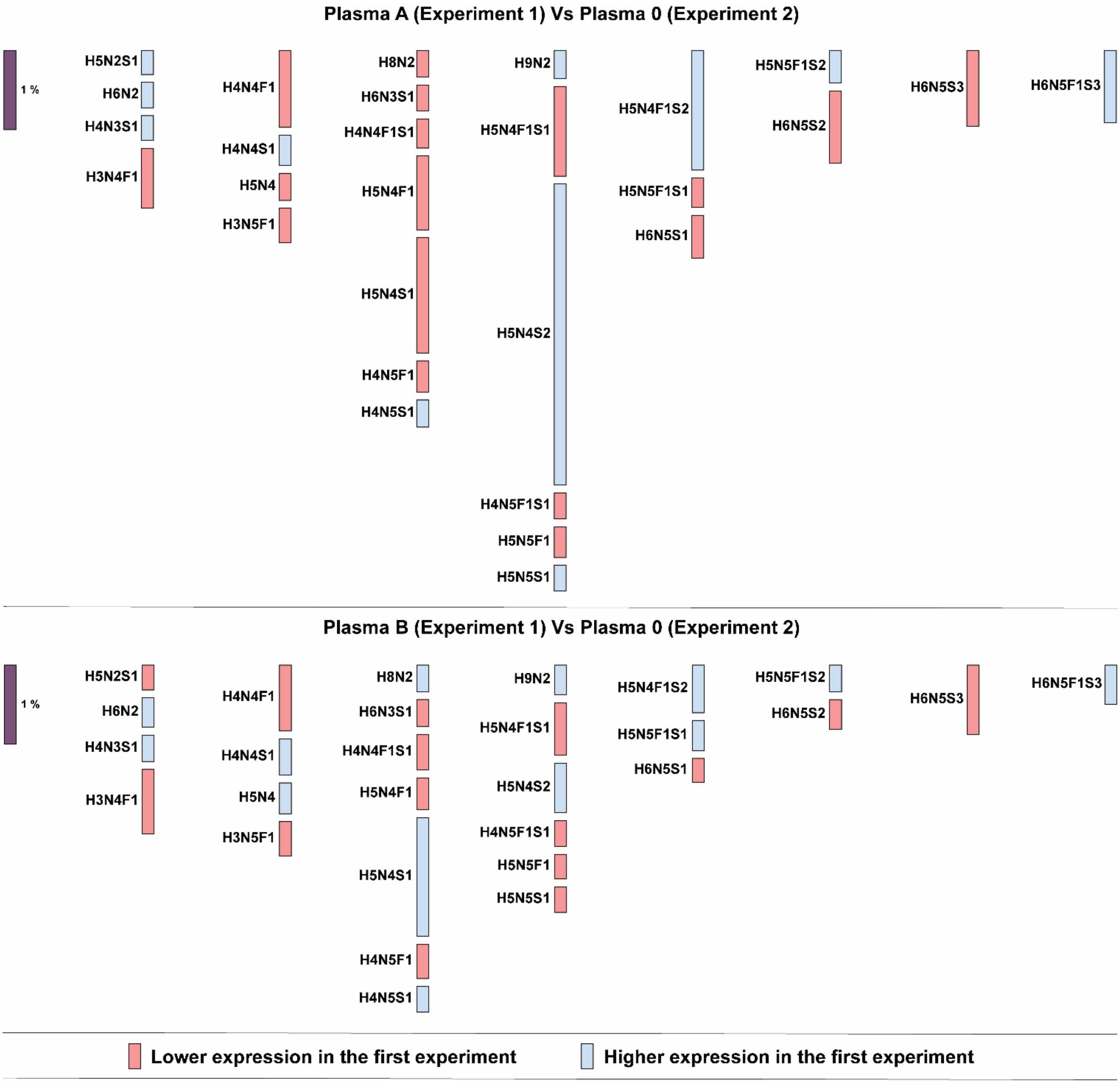


***Figure S12:*** *Differential display of glycan compositions derived from plasma glycoprotein between blood group A vs blood group O and blood group vs blood group O. Compositions are displayed in columns and in each column the overall number of monosaccharides is constant.*

**Source Table S1:** A-F: Pairwise t-test comparison of glycan subclass features among ABO blood group derived from Human plasma.

**A**

| .y. | group1 | group2 | p.adj | p.adj.signif |
|---|---|---|---|---|
| Oligomannose | A | B | 0.0567 | ns |
| Oligomannose | A | O | 0.5040 | ns |
| Oligomannose | B | O | 0.0348 | * |

**B**

| .y. | group1 | group2 | p.adj | p.adj.signif |
|---|---|---|---|---|
| Neutral | A | B | 0.032900 | * |
| Neutral | A | O | 0.000885 | *** |
| Neutral | B | O | 0.006520 | ** |

**C**

| .y. | group1 | group2 | p.adj | p.adj.signif |
|---|---|---|---|---|
| Mono | A | B | 0.000604 | *** |
| Mono | A | O | 0.002290 | ** |
| Mono | B | O | 0.069000 | ns |

**D**

| .y. | group1 | group2 | p.adj | p.adj.signif |
|---|---|---|---|---|
| Di | A | B | 0.000303 | *** |
| Di | A | O | 0.000179 | *** |
| Di | B | O | 0.180000 | ns |

**E**

| .y. | group1 | group2 | p.adj | p.adj.signif |
|---|---|---|---|---|
| Tri | A | B | 0.0542 | ns |
| Tri | A | O | 0.3390 | ns |
| Tri | B | O | 0.1430 | ns |

**F**

| .y. | group1 | group2 | p.adj | p.adj.signif |
|---|---|---|---|---|
| Tetra | A | B | 0.3340 | ns |
| Tetra | A | O | 0.0812 | ns |
| Tetra | B | O | 0.3340 | ns |

**Source Table S2:** A-F: Pairwise t-test comparison of glycan subclass features among ABO blood group derived from Human urine.

**A**

| .y. | group1 | group2 | p.adj | p.adj.signif |
|---|---|---|---|---|
| Oligomannose | A | B | 6.29e−04 | *** |
| Oligomannose | A | O | 1.59e−04 | *** |
| Oligomannose | B | O | 1.15e−05 | **** |

**B**

| .y. | group1 | group2 | p.adj | p.adj.signif |
|---|---|---|---|---|
| Neutral | A | B | 1.23e−03 | ** |
| Neutral | A | O | 1.23e−03 | ** |
| Neutral | B | O | 4.29e−05 | **** |

**C**

| .y. | group1 | group2 | p.adj | p.adj.signif |
|---|---|---|---|---|
| Mono | A | B | 5.59e−05 | **** |
| Mono | A | O | 3.48e−03 | ** |
| Mono | B | O | 1.13e−05 | **** |

**D**

| .y. | group1 | group2 | p.adj | p.adj.signif |
|---|---|---|---|---|
| Di | A | B | 3.41e−05 | **** |
| Di | A | O | 3.41e−05 | **** |
| Di | B | O | 1.09e−06 | **** |

**E**

| .y. | group1 | group2 | p.adj | p.adj.signif |
|---|---|---|---|---|
| Tri | A | B | 0.045300 | * |
| Tri | A | O | 0.000096 | **** |
| Tri | B | O | 0.000277 | *** |

**F**

| .y. | group1 | group2 | p.adj | p.adj.signif |
|---|---|---|---|---|
| Tetra | A | B | 0.3340 | ns |
| Tetra | A | O | 0.0812 | ns |
| Tetra | B | O | 0.3340 | ns |
